# Content-enriched fluorescence lifetime fluctuation spectroscopy to study bio-molecular condensate formation

**DOI:** 10.1101/2023.06.09.544221

**Authors:** Eleonora Perego, Sabrina Zappone, Francesco Castagnetti, Davide Mariani, Erika Vitiello, Jakob Rupert, Elsa Zacco, Gian Gaetano Tartaglia, Irene Bozzoni, Eli Slenders, Giuseppe Vicidomini

## Abstract

Quantitative fluorescence microscopy is experiencing an important revolution thanks to single-photon array detectors. These detectors provide users with so far inaccessible specimen information: The distribution of the specimen’s fluorescence emission at single-photon level and high spatiotemporal sampling. In laser-scanning microscopy, this photon-resolved measurement has enabled robust fluorescence lifetime imaging at sub-diffraction spatial resolution, thus opening new perspectives for structural and functional imaging. Despite these significant advances in imaging, studying the time evolution of biological processes remains a considerable challenge. Here we present a comprehensive framework of live-cell spectroscopy methodologies – compatible with imaging – to investigate bio-molecular processes at various spatiotemporal scales. We use photon-resolved spatial and temporal measurements granted by a single-photon array detector to boost the information content of a unified fluorescence fluctuation spectroscopy and fluorescence lifetime experiment. To demonstrate the potential of this approach, we investigate the phase transition of liquid-like condensates during oxidative stress inside living cells. These condensates are generally found in several cellular processes and exhibit substantial variations in molecular composition, size, and kinetics, posing a significant challenge for quantifying their underlying molecular dynamics. This study demonstrates how the proposed approach reveals the mutual dynamics of different RNA-binding proteins involved in the stress granules formation – inaccessible to imaging alone. We observe condensate formation by performing time-lapse super-resolved imaging of the cellular macro-environment while simultaneously monitoring the molecular mobility, the sub-diffraction environment organization, interactions, and nano-environment properties through fluorescence lifetime fluctuation spectroscopy. We are confident that our framework offers a versatile toolkit for investigating a broad range of bio-molecular processes – not limited to liquid-liquid phase transition – and we anticipate their widespread application in future life-science research.

## Introduction

Fluorescence fluctuation spectroscopy (FFS) is a family of methods that allows quantifying molecular concentration, mobility, interactions, confinements, and aggregations even in crowded and complex environments such as a living cell (1–3). These methods record the fluorescence intensity variations created by fluorescently tagged molecules passing through a small detection volume (or probed region) and statistically analyze these fluctuations to characterize the molecule properties. The most frequently used routine of FFS is fluorescence correlation spectroscopy (FCS), in which the autocorrelation function of the intensity fluctuations is calculated and analyzed in order to grant averaged information about molecular mobility and concentration in the sample.

In laser-scanning microscopy (LSM), FCS uses confocality or two-photon excitation to generate the small (order of femtoliter) detection volume and a fast single-point detector, e.g., a photomultiplier (PMT) or a single-photon avalanche diode (SPAD), to sample the fluorescence signal. However, it has been recently shown that the information content of LSM-based FCS increases by substituting the conventional single-point detector with a fast (≤ *µ*s) array detector able to directly image the probed region (4–6). Indeed, a single-point detector averages out all the possible spatial heterogeneity of the fluorescence signal from the probed region. On the contrary, fast array detectors preserve this spatial information without compromising the temporal sampling. This spatial information has been used to perform several FFS methods in a more straightforward configuration – collectively named comprehensive correlation analysis (CCA) (4) – such as spot-variation fluorescence correlation spectroscopy (svFCS) (7, 8), providing more insights into the investigated bio-molecule diffusive processes. Spot-variation FCS infers sub-resolution spatial heterogeneity within the probed region, by measuring the diffusion times in tunable observation volume sizes – while still measuring the mobility. As a result, svFCS allows distinguishing between free or constrained – caused by sub-diffraction structures – motion, such as trapping from partition domains or hopping due to meshwork.

In conventional svFCS, this was obtained by performing several sequential measurements with either changing the back-aperture objective dimension, thus effectively modifying the hardware of the optical setup, or by applying super-resolution methods (e.g., stimulated emission depletion microscopy) (9–11), with a versatility reduction and a phototoxicity increase. Alternatively, with array detectors different detection volumes are created by integrating the signal, in a post-processing phase, from selected array elements (12, 13). From a single and gentle measurement, it is possible to easily extract the transit time as a function of the effective detection volumes, from which we can quantify both bio-molecule mobility and the type of motion, influenced by the sub-resolution sample organization.

Because our svFCS implementation obtains simultaneously the different transit times, it is able to reveal also fast-scale (in the range of the measurement time, i.e., tens of seconds) temporal heterogeneity – typically lost by conventional svFCS. The first implementation of CCA used a PMT array (i.e., the AiryScan detector) (14), whose analog read-out introduces complex data calibrations in the analysis. Asynchronous read-out SPAD array detectors (15, 16) solved this problem and also opened new perspectives for CCA. Indeed, when combined with dedicated data acquisition (DAQ) systems, asynchronous read-out SPAD array detectors allow for a photon-resolved measurement of the fluorescence signal in the detection volume: Every photon is recorded with a series of spatial and temporal signatures. By sending each SPAD element output to a multi-channel time-tagging DAQ module (Fig.1a), it is possible to tag every single photon with a spatial tag corresponding to the photon arrival position (sub-diffraction precision) within the detector, and a time tag, corresponding to the delay (tens of picoseconds precision) from the experiment start (i.e., absolute time). The amount of information increases, providing the DAQ module with the reference signals from the microscope scanner system and the excitation pulsed laser. Each photon is also tagged with the spatial position of the probed region in the sample and the delay from the fluorophore excitation event (i.e., the photon arrival time). The sample position allows for reconstructing a (raster) image or implementing scanning FCS (sFCS). In sFCS the detection volume is scanned repetitively along a line, a circle, or a raster pattern. In contrast to single-point FCS, scanning FCS allows for investigating different sample positions. Consequently, sFCS reveals large-scale (µm) spatial heterogeneity in the investigated process. Accessing the spatial heterogeneity comes with a price, sFCS is practically limited by the intensity trace sampling (ms in contrast of µs, typical for single-point FCS), thus precluding the detection of fast mobility. Thanks to the temporal precision of SPAD array detector, the photon arrival time allows for integrating the fluorescence lifetime information, i.e., fluorescence lifetime fluctuation spectroscopy (FLFS) and fluorescence lifetime imaging (FLIM).

**Fig. 1.**
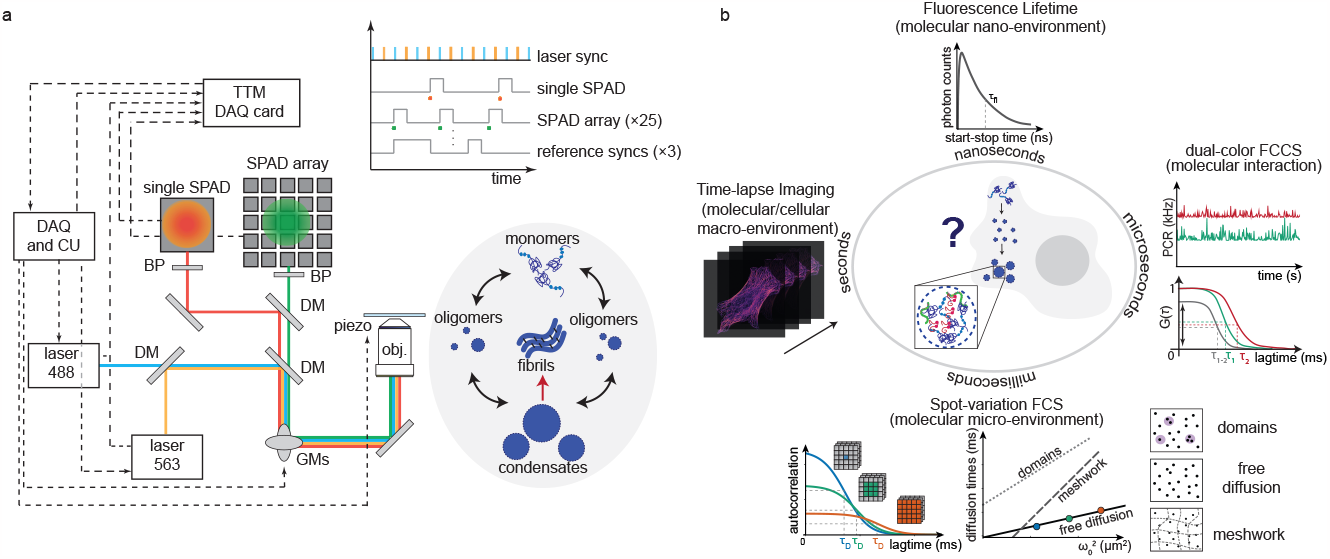
Schematic representation of content-enriched fluorescence lifetime fluctuation spectroscopy. **a** Schematic representation of the optical setup. Two laser beams, to allow for dual-color experiments, are coupled with a dichroic mirror (DM), focused with a 100× objective lens and scanned across the sample with a pair of galvo-mirrors (GMs). The emitted signal collected by the objective lens is de-scanned by the GMs, and split depending on the emitted wavelength and either focused on a SPAD array detector or onto a single-element detector. Every single photon is collected and tagged with a spatial and temporal tag by the BrightEyes-TTM data-acquisition card for time-resolved measurements or by a FPGA-based control unit (DAQ and CU). Our set-up allows for studying both *in vitro* samples, such as purified proteins, and living cells. **b** Thanks to the unique spatiotemporal tags granted by the SPAD array detector, we can fully use the information of every single photon. With the temporal tags, we access several temporal scales. The fluorescence lifetime, on the nanosecond time regime, which gives us information about the chemical nano-environment of single molecules, is measured simultaneously with the microseconds intensity fluorescence fluctuation, giving us information about the measurable diffusion. Granted by the single-photon spatial tags, the diffusion times in the different detected areas can be evaluated (diffusion law), providing information about diffusion modality or the micro-environment of the molecules. These temporal and spatial tags can be provided in a multi-color scheme, allowing us to perform highly informative dual-color cross-correlation measurements.

Recently, we developed an open-source hardware time-tagging module (TTM) – the so-called BrightEyes-TTM – and validated this module in a series of photon-resolved proof-of-principle measurements, both intensity and lifetime-based imaging as well as CCA experiments (17). However, the full potential of our photon-resolved framework has not been yet explored to effectively generate new insights into complex biological processes. Here, we present a comprehensive framework of live-cell spectroscopy and imaging experiments that leverages this massive single-photon information dataset to investigate bio-molecular processes in living cells. Combining the SPAD array detector with the BrightEyes-TTM allows for specimen information across multiple spatial and temporal scales (Fig.1b). The photon arrival time enables calculating the labeled molecule’s fluorescence lifetime, which requires access to the variations of the fluorescence signal at the nanosecond time scale (Fig.1b top). The fluorescence lifetime is known to be a reporter of the nano-environment of the fluorophores (18) (i.e., in the nanometer domain). It has been shown how by using lifetime-sensitive fluorescence probes or FRET construct (Förster resonance energy transfer) that the change in fluorescence lifetime is linked with changes in the biological nano-environment of single molecules (19). Simultaneously, the single-photon absolute time can be binned on the microseconds time scale, creating a time trace of fluorescence fluctuation for each element of the SPAD array detector. These intensity time traces can be analyzed in several ways to extract different data from the same experiment different molecular properties. Here, we focused on two approaches based on FFS methods: svFCS, to simultaneously monitor the molecular mobility and its sub-diffraction environment organization, and dual-color fluorescence cross-correlation spectroscopy (dcFCCS) to monitor interactions between two molecular species (20). In particular, by labeling with two spectrally separable dyes two different types of molecules, the relative amplitude of dcFCCS curves is proportional to the molecular interaction level in the sample (20). A very low amplitude, or a zero cross-correlation, is related to two molecules moving independently while a high amplitude is caused by a coupled movement, indicating an interaction (Fig.1b right). Moreover, since the SPAD array detector is integrated into a LSM, we can perform time-lapse super-resolution imaging by image scanning microscopy (ISM) (21, 22) (Fig.1b left). Therefore, we have a complete platform for high-informative fluorescence experiments where multiple parameters are quantified and simultaneously correlated (i.e., nanoenvironment, sub-diffraction environment, mobility, interaction, and macro-environment) across different spatial and temporal scales.

We showed the potential of our platform by studying the formation of stress granules (SGs) in living cells upon oxidative stress. The formation of SGs is described as a liquid-liquid phase separation (LLPS) event, where membraneless condensates of proteins are formed under specific conditions. The membraneless condensates are often found in different cell cycle stages, helping maintain cellular stability. The condensates are composed of proteins, with typically intrinsically disordered domains, and nucleic acids, e.g., DNA and RNA. They are governed by weak and multivalent interactions between the bio-molecules, and they tend to have spherical shapes, dynamics, and liquid-like properties (23, 24). It is believed that the intracellular components can form reaction compartments by phase separation, forming an environment where multiple biological reactions co-occur (25). SGs are formed mostly by mRNAs and RNA-binding proteins, such as G3BP1 (Ras GTPase-activating protein-binding protein) protein and FUS (FUsed in Sarcoma) protein, and their formation triggers cellular stress response (26). They are known to behave like liquids, as they can assemble and disassemble in response to cellular stimuli. However, their persistent formation or aberration, for example, the inclusion of mutant proteins, has been related to neurodegenerative diseases (27). In particular, some mutations of FUS protein are related to amyotrophic lateral sclerosis (ALS). FUS is a pleiotropic RNA binding protein with several functions in regulating RNA metabolism; although being mainly nuclear, ALS-causing mutations trigger its displacement inside the cytoplasm. This results possibly into a double toxic effect, as it combines a loss of function in the nucleus and a gain of toxic function in the cytoplasm. SGs formation is initiated here by inducing oxidative stress. In its ALS-mutated forms, FUS is also incorporated into the SGs upon stress induction, dysregulating consequently SGs physiology (28). It remains still unclear whether the mislocation and the co-localization of the ALS-mutated FUS inside SGs are causative of ALS or whether this cytopathological aggregation is the response consequence to the stress of degenerated motoneurons. The aggregation of mutated FUS could inhibit the localization of RNAs encoding for proliferating factors, hindering cell recovery after stress and thus increasing neuronal loss. Moreover, the RNA-binding capability of FUS might increase the sequestration of specific mRNAs, which might be essential for neuronal viability.

Protein-protein interaction and protein-RNA interaction are fundamental for SG’s dynamicity and fluidity, thus, it is important to study interactions in combination with the dynamic properties of single proteins. Due to the complexity of the process, only with a multi-parameter platform and on multiple spatiotemporal scales, it will be possible to get a more complete view of the processes behind SG formation, and eventually on SG aberration and aggregation. Despite the specific biological process studied here, the same framework can be applied to several other processes where the dynamics of biomolecules are investigated with multiple physical parameters in unprecedented detail.

## Results

### Validation of Fluorescence Lifetime Fluctuation Spectroscopy

To validate our multi-parameter platform, we started by using test samples. We first measured fluorescent beads dissolved in water. We implemented svFCS with a 5× 5 elements SPAD array detector and a custom laserscanning microscope as explained in (5, 17). We used the photon absolute time to obtain the intensity time trace for each SPAD array element. We integrated the intensity traces to virtually detect the signal from three volumes of observation: The volume from the central element only, the sum of the inner square (Sum 3 ×3), and the sum of all channels (Sum 5× 5) (Fig.2b). We auto-correlated the resulting time traces and fitted the relative auto-correlation functions with a one-component 3D diffusing model (Fig.2a). The diffusion law 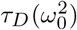 allows distinguishing different modalities of diffusion (5, 8, 29). Here, as the intercept is close to the origin (0.03± 0.03 ms), the diffusion law suggests an expected free diffusing motion (Fig.2c). The measured diffusion coefficient (17.7 ± 0.3) µm^2^/s and the calculated diameter (*r* = 24 nm) correspond well with previous measurements and the manufacturer’s information.

**Fig. 2.**
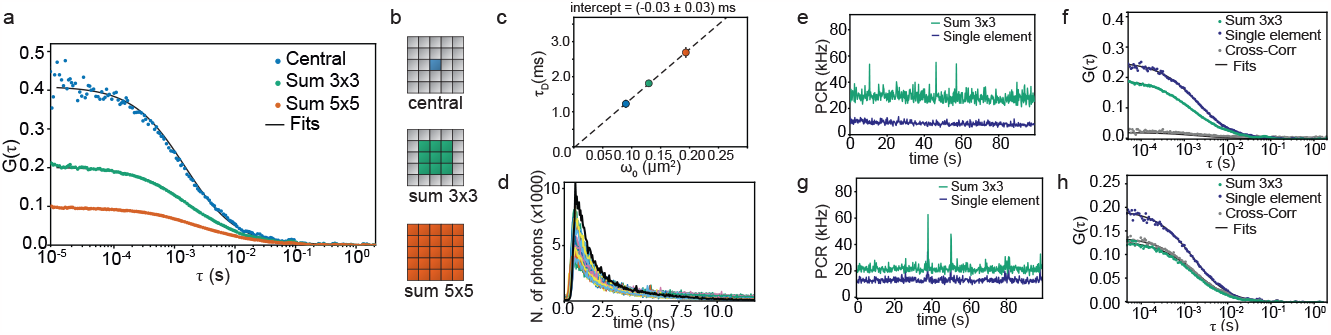
Single measurement of fluorescent nanospheres with the SPAD array detector. **a** FCS measurement of fluorescent nanospheres in water. The three curves correspond to the three volumes considered with the SPAD array detector. The three autocorrelation curves are fitted (black lines) with a model for one diffusing component. **b** Schematic depiction of the area considered on the SPAD array detector for calculating the autocorrelation at the three observation volumes. **c** Spot-variation FCS measurement of fluorescent nanospheres in water. The diffusing times are calculated from the autocorrelation curves shown in **a**. Here, the diffusion law confirms a free motion with an intercept value of (−0.03 ± 0.03) ms and a diffusion coefficient of (17.7 ± 0.3) µm^2^/s, which corresponds to a diameter of 24 nm (20 nm from the manufacture). **d** Histogram of the fluorescence decay calculated from the same dataset used in (a). A histogram of the fluorescence decay is calculated for each of the 25 channels of the SPAD array detector. The black line represents the data from the central detector channel. The fluorescence lifetime value can be retrieved by fitting the data with an exponential function (*τ*_*F*_ = (1.3 ± 0.1) ns). **e-h** Cross-Correlation experiment with fluorescent nanospheres. **e-f** A mix of beads with two colors is used for the measurements (yellow-green and orange-red). The SPAD array detector is equipped with a filter for green fluorescence signals, while the single-element detector with a filter for red fluorescence signals. While the signal is well autocorrelated in every single channel, the cross-correlation shows a low amplitude, suggesting low interaction between the two samples. **g-h** Only yellow-green fluorescent nanospheres are diluted in water and measured. The signal is split equally on the two detector arms, which are equipped with the same detection filter. The intensity time trace can be well-correlated in both detectors (only the Sum 3×3 is shown here for the SPAD array detector). As expected here, the cross-correlation curve has a high amplitude as the signal is the same for both detectors.

We further validate our svFCS approach on a more complex system where mobility is not only restricted to be free diffusion. We investigated the *in-vitro* LLPS of alpha-synuclein protein (*α*-syn). Alpha-synuclein is a pre-synaptic protein found typically in the neuronal brain tissue. Solid-like aggregates of *α*-syn have been identified as one of the main components of Lewy bodies, the pathological neuronal inclusions associated with different neuronal diseases such as Parkinson’s (30). Recently, it has been shown that the aggregation of soluble *α*-syn can be preceded by the phase separation in liquid-like condensates (31). Upon the induction of the LLPS (by mimicking molecular crowding through polyethylene glycol), we followed the condensation process by imaging and svFCS (Fig.S1). We characterize the dynamics of monomeric *α*-syn before the LLPS process. Imaging shows a uniform solution of fluorescent *α*-syn. Likewise, also svFCS reveal only one diffusing component in the solution. As expected, *α*-syn is freely moving in solution with a diffusion coefficient of *D* = (130± 7) µm^2^/s, comparable with previously reported values for monomeric *α*-syn (32). Once *α*-syn proteins start the phase-separation process, round droplet-like condensates are visible by imaging. In this case, svFCS reveals two diffusing components in the solution. One, which corresponds to the dilute phase, and one which corresponds to the condensate phase. Interestingly, we can use the diffusion law 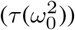 to distinguish the two different types of movement discovered by FCS fitting. The fast component shows a free-diffusion type of motion (intercept *q* close to 0), suggesting that this component is mainly the dilute phase. The slower component shows a positive intercept, indicating a domain-confined diffusion. We disrupted the weak hydrophobic interactions responsible for condensate formation by adding the alcohol 1,6-hexanediol (33, 34).

As expected, upon adding 10% of 1,6-hexanediol we see a clear change in the autocorrelation curve. Even if two diffusing components are still detected, the slower diffusing component shows a lower strength in the condensed phase as well as a lower diffusion coefficient, indicating the starting point of the dissolution of the liquid-like condensates. The experiments on the LLPS of *α*-syn confirm the effectiveness of svFCS on LLPS processes, being able to distinguish the changes in the sub-diffraction environment during the condensation process.

Regarding the original photon-resolved dataset, we calculated the photon-arrival time histograms for each SPAD array element (Fig.2d) on the original fluorescent beads test sample. The photon-arrival time histogram measures the fluorescence decay distribution. The fluorescence lifetime of the tagged molecule is extracted by fitting the decay function. To further investigate potential molecular interactions, we updated the custom microscope with a cross-correlation platform. We obtained this functionality without any change in the data acquisition system and with minimal changes in the optical architecture: We included in the detection path a single-point SPAD connected to the BrightEyes-TTM platform, which can simultaneously acquire signals both from the single-point SPAD and the SPAD array detectors (up to 50 channels in the current version). The fluorescence signal from the detection volume is then split between the two detectors with a 50-50 beam-splitter or a dichroic filter – to separate the intensities spectrally. We tested the system by measuring a mixture of yellow-green and red fluorescent beads. The single-element detector has been equipped with detection filters for the red emission while the SPAD array detector for the green emission (Fig.1a). We can spectrally resolve the fluorescence intensity and separate the signal on the two detectors (Fig.2e). The autocorrelation functions for each color give us a measurement of the diffusion coefficients for each species of beads (Fig.S2). However, the cross-correlation function has a low amplitude, suggesting a shallow interaction between the species. If, with the same microscope configuration, red-only or yellow-green-only beads are measured, one of the two autocorrelation functions is zero (the one from the SPAD detector for red-only beads or the one from the single element detector for yellow-green-only beads) (Fig.S2). Consequently, the cross-correlation also curves have low amplitudes. If, on the other hand, we measured only yellow-green fluorescent beads but with a 50-50 beam-splitter in the detection path and wit green detection filters on the single-element detector, we can retrieve both good autocorrelation curves and a cross-correlation curve with high amplitude.

### Spot-variation FCS reveals protein condensation in living cells

In the context of life sciences, applying these methods allows the simultaneous investigation of function, structure, and temporal evolution of dynamic processes like aggregations, condensations, or interactions. We implemented our methods to investigate the formation of stress granules (SGs) in neuroblastoma-like (SK-N-BE) cells under oxidative stress induction. We focused mainly on Ras GTPase-activating protein-binding protein (G3BP1), a core component of SG assembly network. We measured its interaction with FUsed in Sarcoma (FUS) protein in a WT cell model and in a pathological-related mutated model. In the latter, FUS protein has a single point mutation at the P525L position, which leads to a misplacement of the protein from the nucleus to the cytoplasm, and it is related to amyotrophic lateral sclerosis disease (35, 36). To follow FUS and G3BP1 dynamics, we took advantage of SK-N-BE stable cell lines expressing FUS protein WT or P525L fused to RFP under a Doxycycline inducible promoter (37). Both WT and mutant cell lines were also selected for the stable expression of G3BP1 protein tagged with eGFP under the constitutive Eif1a promoter and the formation of SGs was triggered by applying 0.5 mM sodium-arsenite to the cells.

First, we characterized the sample before applying the oxidative stress. Fig.3a shows a SK-N-BE WT cell uniformly expressing G3BP1-eGFP in the cytoplasm. We performed svFCS to probe its mobility (an example is in Fig.3a top-right) in different cells and different positions of the cell. The autocorrelation curves could be fitted with a one-component 3D diffusing model for the three detection volumes of the SPAD array detector. The diffusion law plot, 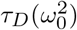 (bottom-right), suggests, in this case, a meshwork-confined type of diffusion. Once the oxidative stress is induced, SGs are visible in cells (Fig.3b), and the correlation curves show a second slower diffusing component in the fits (top-right). The diffusion modality changed compared to before the formation of the stress granules, indicating a change in the micro-environment of G3BP1.

**Fig. 3.**
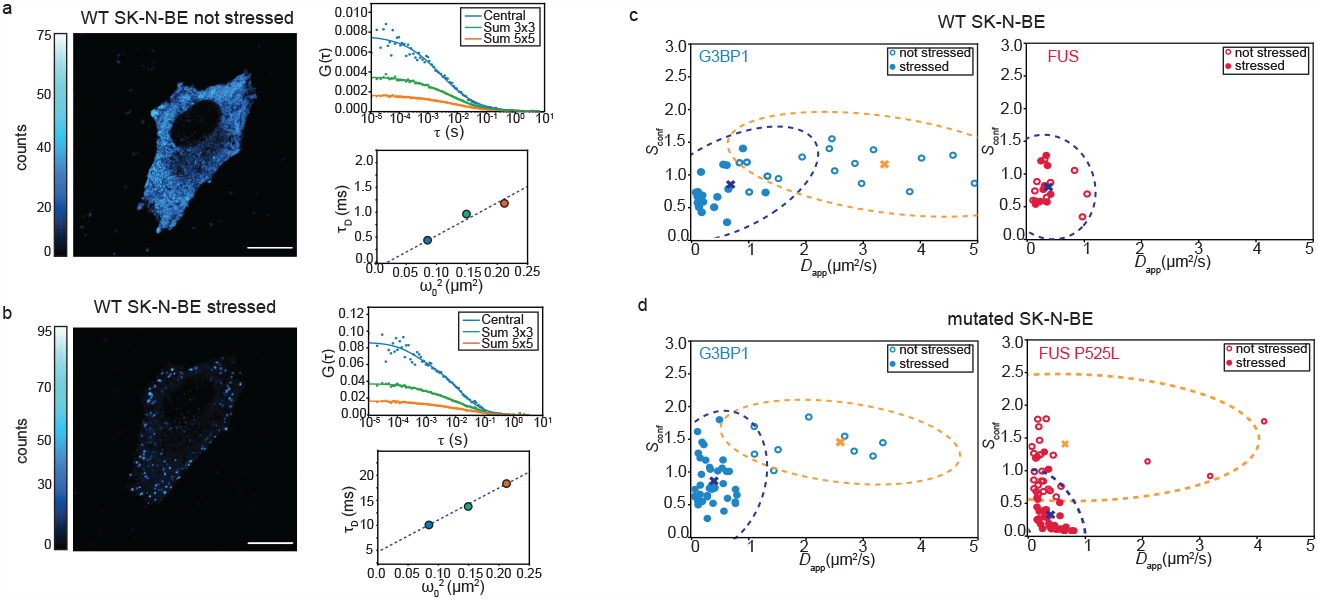
Spot Variation Fluorescence Spectroscopy for LLPS processes in living cells. **a** ISM intensity-based image of an SK-N-BE cell WT expressing G3BP-1-eGFP before oxidative stress. A svFCS measurement is performed in a single point of the cellular cytoplasm by acquiring the fluorescence intensity over time and calculating, off-line, the autocorrelation curves for the three observation volumes (central, Sum 3×3 and Sum 5×5). The diffusing times can be plotted against the area of the observation volume to retrieve the modality of diffusion with the diffusion law. Scale bar 10 µm. **b** ISM intensity-based image of an SK-N-BE cell WT expressing G3BP-1-eGFP during oxidative stress. The SGs are visible in the cell cytoplasm. The change of diffusion/modality of diffusion is reflected in the autocorrelation curves, calculated as in (a). The diffusing times can be plotted against the area of the observation volume to retrieve the modality of diffusion with the diffusion law. Scale bar 10 µm. **c** Confinement strength in relation to the diffusion coefficient of G3BP1 (cyan) and FUS (magenta) protein in WT SK-N-BE cells before the stress (empty circle, *n* = 14 cells measured over 3 independent experiments for G3BP1 and *n* = 6 cells measured over 3 independent experiments for FUS) and after the stress (full circle, *n*= 17 cells measured over 3 independent experiments and *n* = 7 cells measured over 2 independent experiments for FUS). The K-mean clustering algorithm has been applied to the datasets. The crosses represent the centroid position of the found cluster. The dotted ellipses represent the covariance confidence level of each cluster. **d** Confinement strength in relation to the diffusion coefficient of G3BP1 (cyan) and FUS (magenta) protein in mutated SK-N-BE cells before the stress (empty circle, *n* = 8 cells measured over 3 independent experiments for G3BP1 and *n* = 12 cells measured over 3 independent experiments for FUS) and after the stress (full circle, *n* = 20 cells measured over 3 independent experiments for G3BP1 and *n* = 21 cells measured over 3 independent experiments for FUS). The K-mean clustering algorithm has been applied to the datasets. The crosses represent the centroid position of the found cluster. The dotted ellipses represent the covariance confidence level of each cluster.

To simplify this analysis in the case of several measurements, we introduce the confinement strength (*S*_conf_) metric as the ratio between the smallest focal area (central pixel only) and the biggest focal area (Sum 5 ×5) diffusion coefficients. A similar metric was already used to represent how strong molecules are confined in domains while diffusing (38).

When *S*_conf_ is 1 the diffusion coefficients are not dependent on the detection volumes, which is the case when molecules are freely diffusing. If *S*_conf_ is bigger than 1 the molecules are moving in a dense meshwork, while when *S*_conf_ is smaller than one the molecules are showing a confined-domain type of movement. The confinement strength allows us to parameterize the movement of the investigated molecules and compare the different cellular conditions. In the WT condition, G3BP1 proteins restrict their mobility in the oxidative stress condition (Fig.3c left). The apparent diffusion coefficients (in the *x*-axis, empty symbols: pre-stress, filled symbols: stress granules), i.e. the ones measured with the whole detector areas, are smaller after the SGs are formed. In addition, after the stress, the confinement strength is lower than one, indicating a domain/confined behavior. Contrarily, FUS proteins, in the WT cell condition, do not seem to change behavior or mobility (Fig.3c right).

In the pathologically mutated cellular condition (P525L), where FUS protein is misplaced in the cell cytoplasm, G3BP1 has a similar behavior compared to the WT condition. The diffusion coefficients of G3BP1 decrease in the stress condition, as well as its confinement strength which moves towards the confinement-domain region. In opposition to the WT cell condition, FUS shows here different types of mobility before and after the stress. This suggests a change in the interaction or in the type of motility in FUS proteins. While their diffusion velocity does not seem to change in the stress granule formation process, the way that FUS proteins move is different.

In both WT and mutated SK-N-BE cell conditions, clear differences of diffusion coefficients of G3BP1 before (*D*_WT_ = (2.3 ± 1.2) µm^2^/s, *D*_mut_ = (2.1± 0.8) µm^2^/s) and after the stress (*D*_WT_ = (0.4 ±0.3) µm^2^/s, *D*_mut_ = (0.3±0.2) µm^2^/s) are measured, indicating a strong reduction in the protein mobility during the formation of the SGs (Fig.S3 left). The confinement strength of G3BP1 shows a more complicated behavior, reflecting the several roles that G3BP1 has inside the SGs (Fig.S4 left). In the WT cell condition, before the stress, G3BP1 is diffusing freely in the cytoplasm (*S*_conf_ = 1.1 ±0.2), while once the SGs are assembled, the confinement strength is lower than one (*S*_conf_ = 0.7± 0.2), indicating a confinement behavior in the granules. In the mutated cell condition, G3BP1 seems to move initially in a meshwork (*S*_conf_ = 1.4 ±0.2), while, upon the formation of SGs its mobility is confined in domains (*S*_conf_ = (0.8± 0.3)), confirming the formation of liquid condensates.

Regarding FUS protein in both WT and mutated cell condition, we did not observe a change in the diffusion coefficients before (*D*_WT_ = (0.5± 0.3) µm^2^/s, *D*_mut_ = (0.2 ±0.1) µm^2^/s) and after (*D*_WT_ = (0.30± 0.08) µm^2^/s, *D*_mut_ = (0.4± 0.2) µm^2^/s) the formation of the SGs (Fig.S3 right). In fact, FUS protein is already partially segregated into condensates before the induction of oxidative stress (Fig.5 a). However, the two cell conditions differ regarding the confinement strength, thus in the type of mobility of FUS protein. In the WT cell condition, as expected, we do not observe differences in the confinement strength in not-stressed and stressed conditions.

We measured before the stress a value for the confinement strength of 0.8 ±0.2, and during the stress a very similar confinement strength value of 0.9± 0.3. It is important to notice that FUS in the WT cell condition is found only in the nuclei and it is not involved in the formation of SGs in the cytoplasm.

On the contrary, in the mutated cell condition where FUS is found in the cytoplasm, the confinement strength significantly decreases (from an average value of 1.0 ±0.4 before the stress, to an average of 0.3±0.3 in the stress granules) upon the formation of the SGs (Fig.S4 right), indicating a confinement-type of mobility. The large standard deviations in the values of the confinement strength for FUS protein are related to the biological variability that is encountered inside SGs. FUS proteins interact with multiple partners inside SGs, from other RNA-binding proteins to RNA molecules.

We performed a k-mean clustering algorithm that partitions the dataset in *k* clusters, minimizing each observation’s distance to the nearest mean. We performed the elbow’s method to retrieve the number of clusters *k* for the analysis. In almost all cases, we found that 2 clusters were enough to describe our datasets. In all cases, the clustering algorithm can classify the data points into two groups, which correspond well to the pre-stress and post-stress conditions. To visually assess the clustering algorithm, we calculated the covariance confidence level based on Pearson’s coefficient. We plotted its values on the clustered data (the dotted ellipses in Fig.3c-d).

We followed further the process of SG formation by investigating the recovery of the cells from the stress condition over time. We followed the changes on the cellular and sub-cellular level during the stress retrieval by ISM time-lapse imaging (Fig.S5a). The SGs are dissolved over the course of 5 hours. This proves the reversibility of the induced stress, confirming that the investigated process is a liquid-liquid phase separation. In between the ISM imaging, we performed single-point spectroscopy measurements in multiple positions of the cells. In this way, we have a time-lapse measurement also of the mobility of the single proteins. The apparent diffusion coefficient and the confinement strength can be correlated with the kinetics of the recovery process. In Fig.S5b the data points are color-coded with the recovery time. The clustering algorithm shows a similar clusterization as for the stress-inducing process (Fig.3c-d). It’s important to note that we are able to not only demonstrate the dissolution of the SGs with imaging but also directly on a single protein level, well below the optical resolution.

### Fluorescence lifetime reveals protein confinement

Thanks to the photon-arrival time granted by the coordination between the BrightEyes-TTM and the SPAD array detector, we can analyze the acquired data to build the photon arrival-time (or decay) histogram of the fluorescence signal and perform time-resolved spectroscopy. In the context of imaging, we demonstrated the potentiality of the SPAD array detector combined with the BrightEyes-TTM platform (17), by implementing fluorescence lifetime image scanning microscopy (FLISM). Briefly, first, we used the adaptive pixel-reassignment (APR) algorithm to reconstruct the high spatial resolution and high signal-to-noise ratio, image-scanning microscopy (ISM) intensity image (21) (Fig.4a bottom-left, G3BP1-eGFP expressing cells under oxidative stress). Secondly, we quantify the fluorescence lifetime by both fitting the photon arrival-time histograms for each pixel with a single-component exponential decay function (Fig.4a-b top-right) and by phasor analysis (Fig.4c-d). Monitoring the changes in the fluorescence lifetime is crucial as it can be related to structural or functional changes in the cellular structure. Here, once the cells are under oxidative stress and the SGs are formed, we measured a significant (t-test, p-value = 2e-6) decrease in fluorescence lifetime of G3BP1-eGFP inside the SGs (Fig.4f). The segmentation based on the fluorescence lifetime signal of G3BP1-eGFP allows real-time tracking of the changes in the protein environment (structure but also function), i.e., the formation of SGs. By a simple centroid-based segmentation based on the phasor plot, and consequently, on the fluorescence lifetime value, the ISM intensity image allows highlighting the difference between long and short lifetime (Fig.4d). In our case, we are able to automatically segment the SG position within single cells, without any intensity-based threshold assumption or fitting procedure.

**Fig. 4.**
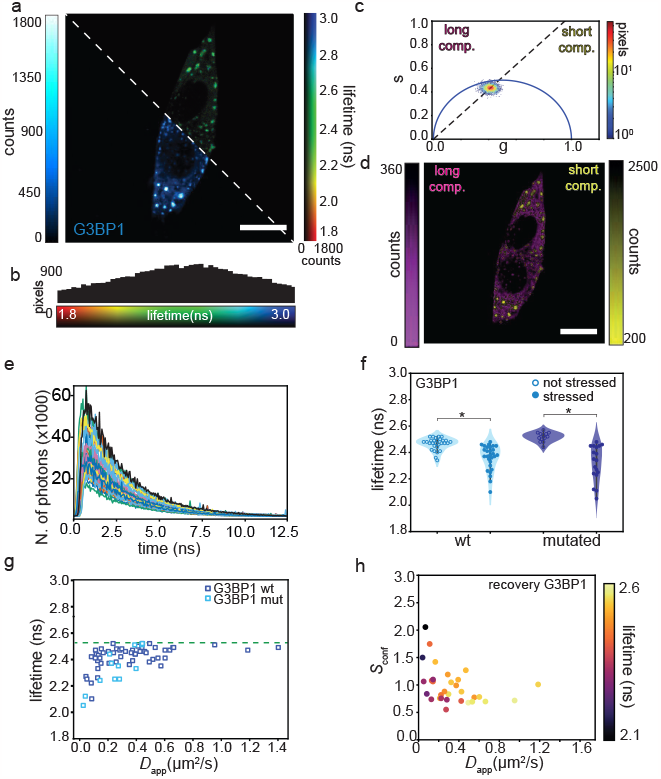
Fluorescence lifetime Fluctuation Spectroscopy. **a** SK-N-BE WT cell expressing G3BP1-eGFP under oxidative stress. Bottom-left ISM reconstructed intensity-based image, top-right ISM reconstructed fluorescence lifetime-based image. Scale bar 10 µm. **b** Histogram of the fluorescence lifetime values of the image in (a). **c** Pixel intensity thresholded phasor plots. Number of pixels versus the polar coordinate (10% thresholds) of the image shown in (a). **d** Phasor-based segmentation. ISM intensity images were obtained by back projecting the points on the left of the phasor centroid (dotted line), representing pixels of the image with a long lifetime, and the points on the right of the phasor centroid, representing pixels of the image with a short lifetime. Scale bar 10 µm. **e** Time decay histograms for the different SPAD array detector pixels, central pixel in black. **f** Violin plots of G3BP1 fluorescence lifetime in the diluted cytosolic phase (on the left) and in the stress granule phase (on the right). The blue circles are measurements of the mutated cell condition (*n* = 29 for the not-stress condition, *n* = 28 for the stress condition) while the cyan squares are measurements in the WT condition(*n* = 13 for the not-stress condition, *n* = 18 for the stress condition). In both cases, the fluorescence lifetimes are significantly different when measured in the diluted or in the condensed phase (p-value < 0.05). **g** Fluorescence lifetime of G3BP1 in WT cells (dark blue, *n* = 20) and in the mutated cells (cyan, *n* = 25) correlated with the apparent diffusion coefficient. The dotted line corresponds to the fluorescence lifetime of eGFP molecules. **h** Fluorescence lifetime of G3BP1 in mutated cells correlated with the confinement strength and the diffusion coefficient during the recovery from the induced stress (*n*= 31).

**Fig. 5.**
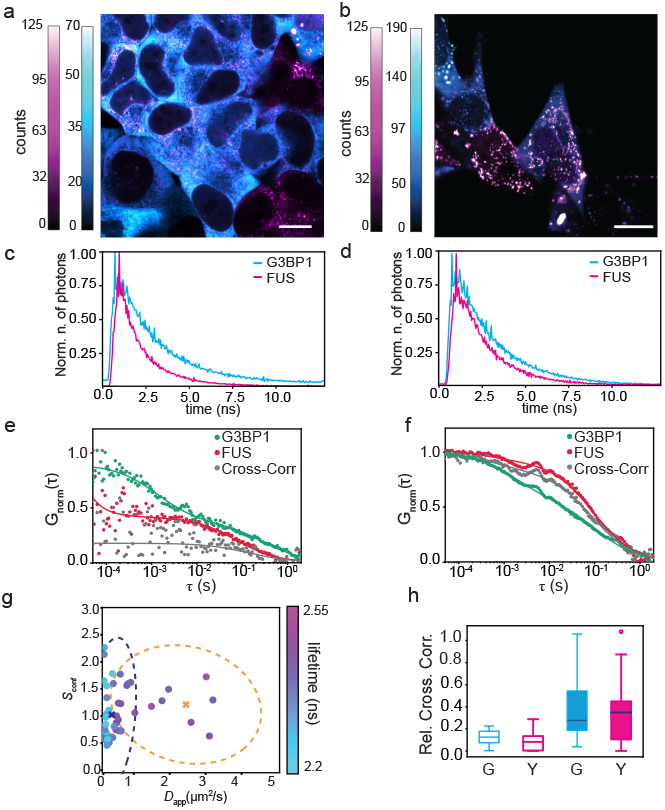
Cross Correlation Fluorescence lifetime Spectroscopy. **a** Intensity-based ISM image of an SK-N-BE mutated cell expressing G3BP1-eGFP (cytoplasm, in cyan) and FUS-RFP (cytoplasm, magenta) before the oxidative stress. Scale bar 7 µm. **b** Intensity-based ISM image of a SK-N-BE mutated cell expressing G3BP1-eGFP (cytoplasm, in cyan) and FUS-RFP (cytoplasm, magenta) after the oxidative stress. Scale bar 5 µm. **c** Time decay histograms for G3BP1 (cyan) and FUS (magenta) in SK-N-BE mutates cells (a) before the oxidative stress. **d** Time decay histograms for G3BP1 (cyan) and FUS (magenta) in SK-N-BE mutates cells (d) during the oxidative stress. **e** Autocorrelations and cross-correlation curve (grey) calculated with the same data shown in c for G3BP1-eGFP (green), FUS-RFP (red), in the cytoplasm before the SGs formation. **f** Autocorrelations and cross-correlation curve (grey) calculated with the same data shown in d for G3BP1-eGFP (green), FUS-RFP (red) inside a SG. **g** Confinement strength and diffusion coefficient of G3BP1-eGFP in mutated SK-N-BE cells in relation to their fluorescence lifetime. The K-mean clustering algorithm has been applied to the datasets. The crosses represent the centroid position of each cluster. The dotted ellipses represent the covariance confidence level of each cluster (*n* = 32). **h** Relative cross-correlation amplitude. Empty columns represent measurements on SK-N-BE mutated cells before the stress. Filled columns represent measurements of SK-N-BE mutated cells after the stress. Cyan is the relative amplitude in relation to the G3BP1 signal while the magenta is in relation to FUS. The horizontal line in each box represents the median, box edges are the 25th and 75th percentile, and the vertical line extends to the minimum and maximum data points.

In the context of FFS, we showed here how we could enhance the information provided by the imaging experiments by performing FLFS during the SG formation. By binning in time the photon arrival time on the microsecond time scale, we are able to perform svFCS on the very same position and time point of fluorescence lifetime measurements. By correlating the intensity in time on the three SPAD array detector volumes and, simultaneously, employing the photon arrival-time to create the time-decay histogram (Fig.4e), we are able to correlate the information about dynamics and sub-diffraction changes with the changes in the nano-environment of fluorescent-tagged molecules (Fig.4g), granting us the link between structure and mobility. In our case, the fluorescence lifetime decreases in correlation with the diffusion coefficient for G3BP1 both in the WT and in the mutated cell conditions, indicating a rearrangement of the dye molecules during the SG formation. A decrease in fluorescence lifetime has already been reported during amyloid fibril formation (39). The fluorescence lifetime reaches a plateau at a value around 2.55 ns which corresponds well to the fluorescence lifetime value of eGFP (between 2.4 and 2.6 ns (40)), suggesting that there is no influence on the fluorescence lifetime before oxidative stress when G3BP1 is diffusing almost freely in the cytoplasm. The same clusterization found for the confinement strength (Fig.3c-d) is furthermore validated by the life-time information, as it is also clustered, finally correlating the fluorescence lifetime changes with the changes in protein dynamics (Fig.4h) even when dissolving the SGs during stress recovery.

### Dual-color FCCS reveals proteins interaction

We finally performed two-color cross-correlation spectroscopy (dcFCCS) measurements to demonstrate the interaction between G3BP1 and FUS during SG formation. In this case, we used the same microscope configuration as in the test-beads experiments: G3BP1-eGFP signal is acquired with the SPAD array detector and FUS-RFP signal with the single-element detector. We performed the ISM imaging for G3BP1 protein, while conventional confocal imaging for FUS protein. The two detection volumes were calibrated with fluorescent beads measurements (Fig.S6). The detection volume of the single-element detector corresponds in dimension with the detection volume of the inner 3× 3 elements of the SPAD array detector (about 330 nm). Before every dcFCCS, the sample was visually inspected, and ISM or confocal images were acquired. In the WT cell condition, the cross-correlation measurements are not well performing as the signal of one of the proteins, depending on the measured position, is basically noise-dominated due to the different compartmentalization of the two proteins (Fig. S7). While in the SK-N-BE cells with the mutation, both FUS and G3BP1 are co-present in the cell cytoplasm already before the stress condition (Fig.5a). Adding sodium-arsenite stress is induced and SGs are formed in the cell cytoplasm. The SGs are colocalizing with the condensation of FUS (Fig.5b). In the pathologically-mutated SK-N-BE condition, we are able to record the fluorescence signal from both proteins simultaneously and reconstruct the intensity time trace for both detectors, as well as the photon arrival-time histogram (Fig.5c-d). Autocorrelation and cross-correlation of G3BP1 and FUS before and during stress are shown in Fig.5(e-f). The amplitude of the normalized cross-correlation function is null in the case of WT SK-N-BE cells (where FUS and G3BP1 are not colocalized), while it is lower before the stress in comparison to inside the SGs. Before the formation of the SGs, G3BP1 and FUS both show good autocorrelation curves. Surprisingly, the cross-correlation curve still can be calculated and fitted, indicating some degree of interaction between the two proteins on longer time scales. This might suggest the formation of few small oligomers between G3BP1 and mutated-FUS protein. Measurements inside SGs (Fig.5f) show longer time decay on both autocorrelation curves and on the cross-correlation curve, indicating the formation of bigger condensates containing both proteins. The amplitude of the cross-correlation function is higher (grey curve in Fig.5f) compared to before the stress induction, suggesting not only the formation of larger condensates but also the increase in interaction strength between the two proteins. This is also well represented by the relative amplitude of the cross-correlation curves (Fig.5h) before (around 0.1) and after the formation of the SGs (around 0.4). The measured increase indicates a strong interaction of G3BP1 and FUS after the formation of the SGs.

We can furthermore include information about the protein nano-environment, taking into account the fluorescence life-time changes in the sample. We can finally combine, in a single dataset, the cross-correlation analysis with the spot-variation analysis and the fluorescence lifetime information, gathering a complete highly-informative group of information for studying SG condensation. Similarly to the experiments on confinement strength on G3BP1, the data shows a similar clustering (Fig.5g, data clustered with k-mean algorithm), and two clusters were found.

## Discussion

This study presents a comprehensive fluorescence-based approach to investigate biomolecular processes in living cells. Our method combines a conventional fluorescence laser-scanning microscope, an asynchronous read-out SPAD array detector, and a time-tagging DAQ module to implement single photon-resolved measurements of the specimen’s fluorescence signal. These measurements reveal the sample information across a wide range of spatiotemporal scales – from nanoseconds to seconds, from nanometers to micrometers. Remarkably, our single-photon fluorescence-based platform allows for high-resolution structural and functional imaging through time-lapse FLISM. Moreover, the platform integrates within a single measurement fluorescence lifetime analysis, dcFCCS, and svFCS. For a given fluorescent-tagged biomolecule, this multi-scale and multi-parameter dataset quantifies the molecule’s macro-environment (or sub-cellular environment), nano-environment, dynamics, sub-diffraction environment (or mobility mode), and interactions while performing its duty during biological processes.

We show the potential of our method investigating the formation of stress granules (SG) through liquid-liquid-phase separation (LLPS). Although conventional and super-resolved imaging techniques can provide valuable insights, they are insufficient to elucidate dynamics at the single-molecule level and, at the same time, to measure changes in molecular interactions or the properties of the nano-environment. To address these limitations, we showcased the capabilities of our integrated platform in quantifying the dynamics of specific RNA-binding proteins, such as G3BP1 and FUS, behind the formation of SGs in SK-N-BE live cells. We proved the viability of our method on both WT cells and mutated cells – that carry one of the typical protein mutations of ALS patients.

While imaging and conventional FCS can reveal only partial information about the role of G3BP1 in the SG formation, our integrated approach provides novel insights. Intensity-based imaging easily captures the increase and decrease of local concentration of G3BP1 during the three phases of the SG formation – pre-stress, post-stress, and recovery. Concurrently, conventional FCS shows changes in the diffusion coefficients of G3BP1 during the SG assembly and dissolution. Nonetheless, it is only through svFCS and confinement strength analysis that we can validate the dynamic partition behaviour of G3BP1, thereby confirming the liquid-condensate nature of the SGs. We observed similar dynamics of G3BP1 both in WT and mutated cells. The advantages of our comprehensive fluorescence-based approach over conventional methods become even more evident when we study the FUS dynamics. Although conventional FCS can reveal only the change in the diffusion coefficients during the three phases of SG formation, it is unable to fully capture alterations in the dynamics of FUS proteins in mutated cells upon stress induction. In stark contrast, the proposed svFCS reveals a significant difference in the confinement strength of the mutated FUS protein before and after the stress induction, indicating a difference in the bio-molecular process underneath. Upon comparing the results for G3BP1 and FUS, we observe that the two proteins exhibit distinct dynamics before stress induction. However, once the two proteins are internalized into the SGs, their diffusion coefficients and confinement strengths become strikingly similar, hinting towards an interaction process. We directly confirm this interaction by measuring the cross-correlation function between FUS and G3BP1 during the stress induction. The amplitudes of the cross-correlation functions increase in measurements inside the SGs, signifying a clear interaction between mutated FUS protein and G3BP1.

In addition to the insights provided by fluorescence fluctuation spectroscopy and intensity-based imaging, our approach combines these findings with the fluorescence lifetime analysis. Recently, the fluorescence lifetime has garnered the interest from various groups, being used as a reporter for functional changes inside living samples (19). We observe changes in the fluorescence lifetime due to the protein compartmentalization after the formation of the SGs. We expect the fluorescence lifetime to be of great importance in studying protein interaction and structural changes, complementing its intrinsic metabolic information, by using it to perform, for example, FLIM-FRET experiment. Once again, our integrated platform offers a simpler and more gentle approach for comprehensive measurement of biological samples during kinetic processes.

The clear dual interaction that we confirm between mutated FUS and G3BP1 inside the SGs might be driven by RNA molecules, such as long non-coding RNAs (lncRNAs) (41– 43), which are known to be present inside the SGs. In perspective, we envision our methods playing a crucial role in uncovering the primary involvement of RNA molecules in various neurodegenerative diseases characterized by the formation of insoluble aggregates, such as Alzheimer’s and Parkinson’s diseases. In general, our platform will be pivotal in directly monitoring the movement and the dynamics of single RNA molecules in the context of RNA and condensate therapeutics (44).

Since we foresee a significant shift from conventional single-element detectors to SPAD array detectors in fluorescence laser-scanning microscopy, we expect the proposed approach to be widely adopted for understanding LLPS and elucidating all biomolecular mechanisms underlying the most important cell functions. Due to the inherent variability of cellular processes, we firmly believe that only a comprehensive approach, where multiple quantitative methods are combined, like the one presented here, will effectively help investigate the kinetics and mechanism behind them. We know that a dominant technology, such as the SPAD array detectors, which enables access to an extensive single-photon dataset will allow for numerous additional analyses.

We anticipate that our platform will enhance the information obtained during fluorescence spectroscopy and imaging measurements, being fully compatible with any type of labeling, sample preparation, or biological process. Similar to how super-resolution microscopy has revolutionized structural imaging, we expect our method to transform the investigation of kinetic processes in living cells. Our comprehensive fluorescence-based platform empowers researchers to gain a deeper understanding of complex biological processes and molecular interactions at multiple scales.

With this vision in mind, we are contributing to this revolution by sharing all the data analysis software presented in this work. This software will be integrated into a more general experimental environment, which also includes our BrightEyes-TTM hardware (17), recently published by us as an open-hardware system for time-resolved measurements with SPAD array detectors.

## Methods

### Laser-scanning microscope

#### Optical architectures

To perform all the experiments, we modified a custom laser-scanning microscope designed for fluorescence fluctuation spectroscopy and imaging with SPAD array detectors (5, 6, 17). Specifically, the microscope uses a 5× 5 SPAD array fabricated with a 0.16 µm BCD technology (16) and cooled to -15^*◦*^C to reduce the dark-noise (6). We additionally integrated a single-element SPAD detector ($PD-050-CTC-FC, Micro Photon Devices, Italy) to register simultaneously the fluorescence light within two different spectral windows, i.e.,to perform dual-color experiments.

Briefly, the microscope excites the sample with two triggerable 485 nm and 561 nm pulsed laser diodes (LDH-D-C-485 and LDH-D-TA-560B, PicoQuant, Germany). We combined the two beams with a dichroic mirror (F43-088, AHF Analysentechnik, Germany) and coupled them into a polarised maintaining fiber before sending them to the microscope. The lasers are controlled by a pair of laser drivers which can be synchronized via an oscillator module (PDL 828-L “SEPIA II”, Picoquant). The oscillator also provides the synchronization signal – synchronized with the excitation pulses – to feed into the time-tagging module and to measure the photon-arrival times. The two co-aligned and collimated excitation beams are deflected by a pair of galvo-mirrors – to scan the probed region across the specimen – and focused on the sample by using a 100 ×/1.4 numerical aperture objective lens (Leica Microsystems, Mannheim, Germany). The emitted fluorescence signal is collected by the same objective lens, de-scanned by the two galvo-mirrors, and focused on the SPAD array detector or on the core of the single-mode fiber of the single-element SPAD detector. To spectrally separate the fluorescence between the two detectors, a short pass filter (F38-534, AHF Analysentechnik) is introduced in the optical path before the two detectors.

### Control and data-acquisition system

We managed the microscope operations with an FPGA-based control unit (PRISM-Light control unit, Genoa Instruments, Genoa, Italy) and the BrightEyes-MCS microscope control software. The microscope control unit drives the galvo-mirrors and the axial piezo-stage to position the probed region on different sample locations or scan the probed region across the whole sample. The control unit also records the signals from the SPAD (array and single-element detectors) in synchronization with the scanning beam system and provides – as outputs – the relative reference signals (i.e., pixel clock, line clock, and frame clock). The BrightEyes-MCS software is based on LabVIEW (National Instruments, Austin, TX) and Python and uses as a backbone the Carma application (Genoa Instruments, Genoa, Italy) (45). The BrightEyes-MCS provides a graphical user interface to control the microscope acquisition parameters (e.g., scanned region, pixel size, axial position, pixel dwell time), to register the digital signals of the SPADs, and to visualize the recorded signals (e.g., intensity images, time traces, and correlations). In the context of fluorescence fluctuation spectroscopy, the BrightEyes-MCS software allows recording the fluorescence signal from each SPAD with a sampling (temporal bin) down to 500 ns. We used this acquisition modality to perform all experiments in which the fluorescence lifetime information is discarded: The photons registered by each SPAD are counted within the temporal bin to obtain the different intensity time traces. When we performed fluorescence lifetime analysis in combination with imaging or spectroscopy, we interleaved the BrightEyes-TTM card between the detectors and the control unit (17): We connected all the SPAD outputs – transistor–transistor logic (TTL) signals – to the time-tagging module, together with the synchronization signal from the laser-driver and the reference signals from the scanning beam systems. The time-tagging module also replicates the input signals of the 3×3 central elements of the SPAD array detector and sends them to the control unit of the microscope. To differentiate the two data acquisition modalities, we named the first intensity-based measurement, i.e., we do not use the BrightEyes-TTM, and the second lifetime-based measurement, i.e., we register the photon-arrival times with the BrightEyes-TTM.

### Samples preparation

#### Calibration samples

For calibrating the confocal volumes of our system, we used a solution of YG carboxylate fluoSpheres (REF F8787, 2% solids, 20 nm diameter, actual size 27 nm, exc./em. 505/515 nm, Invitrogen, ThermoFisher,Waltham, MA, USA) diluted ×5000 in ultrapure water or a solution of Red carboxylate fluoSpheres (REF F8786, 2% solids, 20 nm diameter, exc./em. 580/605 nm, Invitrogen, ThermoFisher) diluted 3000 ×in ultrapure water.

#### Alpha-Synuclein Purification and Labeling

*Wild type α*-syn and *α*-syn with the addition C141 (*α*-syn^C141^) were respectively purified as described in (46). Briefly, both proteins were recombinantly produced in *E. coli*. The pT7-7 aSyn C141 plasmid was a gift from Gabriele Kaminski Schierle (Addgene plasmid #108866; RRID: Addgene_108866). After lysis with boiling, ammonium sulphate precipitation, and dialysis, the proteins were subsequently purified via anion-exchange and size-exclusion chromatography with Hi-Trap™Q and HiLoad™16/600 Superdex™75 columns (Cytiva, USA). The cysteine in position 141 was used to label *α*-syn^C141^ with Atto-488 via a maleimide reaction according to the manufacturer’s instructions. Briefly, the dye is dissolved into water-free DMSO (Thermo Fisher) to a concentration of 10 mM. *α*-syn^C141^ dissolved in 20 mM phosphate buffer was incubated with 1.3 molar excess of Atto-488-maleimide (ATTO-TEC GmbH, Germany) for 120 minutes at room temperature. After the incubation, the labeled protein is separated from the free Atto-488 molecules via PD-10 desalting columns (Cytiva, USA). The tagged protein was immediately checked for concentration, aliquoted, frozen, and kept at -80°C. *α*-syn and free dye concentrations are measured by UV-vis absorption spectroscopy (Nanodrop ND-1000, Thermo Scientific Technologies).

#### Cell culture

Human neuroblastoma cell line (SK-N-BE) stored at -80°C in freezing medium (RPMI Gibco® supplemented with 20% FBS Gibco and 10% DMSO), were quickly defrosted at 37°C, using a thermostatic bath. After removing the freezing medium by centrifugation (5 min at 1000 rpm), cells were resuspended in the appropriate amount of maintaining medium (RPMI Gibco® supplemented with 10% FBS, 2 mM GlutaMAX Sigma®, 1% Penicillin/Streptomycin Sigma®, 1 mM Sodium Pyruvate Gibco) and plated to allow maintenance and amplification. For FUSRFP measurements, its expression was induced with 24 hours administration of 50 ng/ml Doxycycline.

#### Plasmid construction and stable cell lines selection

The transposable element vectors for inducible expression of RFP-FUS^*wt*^ and RFP–FUS^*P* 525*L*+^ were described in (37). The plasmid for G3BP1-eGFP expression was cloned starting from an ePB-bsd-Eif1a (PiggyBac transposable vector) backbone (47) using sequential In-Fusion Cloning (Takara Bio Inc., Otsu, Japan) reactions. CloneAmp HiFi PCR Premix (Clontech) was used for all the PCR amplifications. To produce SK-N-BE cell lines stabling expressing G3BP1-eGFP, 2.5 × 10^5^ cells were plated on 6 cm dishes. The next day cells were transfected with a solution of Optimem (Thermo Fisher), 5 µg of specific plasmid, 0.5 µg of hybrid transposase plasmid and Lipofectamine 2000 transfection reagent (Invitrogen) with a 1:2.5 DNA to transfection reagent ratio.

### FLISM and FLFS experimental protocol and analysis

For the experiments with fluoSpheres a droplet of fluoSpheres diluted in ultrapure water was poured on a coverslip for the FLFS measurements; a fresh sample solution was prepared for each measurement. All measurements were performed at room temperature. For the experiments with *α*-syn, the protein was previously filtered with 0.22 *µ*M Millex Syringe filters (Durapore PVDF, Merck KGaA, Germany) in order to exclude the presence of potential protein oligomers or aggregates. For the monomeric *α*-syn in non-LLPS condition, 700 nM Atto488-maleimide *α*-syn^C141^) was diluted in 20 mM phosphate buffer and 100 mM KCl (Sigma-Aldrich); for the LLPS experiment, Atto488-maleimide *α*-syn^C141^) and *wild-type α*-syn^C141^) were mixed with a ratio of 1:20 in the LLPS buffer (20% poly-ethylene glycol 8000 (PEG-8K, P1458, Sigma-Aldrich in 20 mM phosphate buffer. All measurements were performed at room temperature by dropping 100 *µ*l in a *µ*-Slide 8 well (Ibidi GmBH, Germany). For disruption of *α*-syn LLPS, after 30 minutes LLPS was triggered, 1,3-Hexanediol (Sigma-Aldrich), previously diluted in LLPS buffer, was added to the sample with a final concentration of 10%. For the experiments with SK-N-BE cells, cells were seeded onto a µ-Slide 8 well plate (Ibidi GmbH) and imaged in Live-Cell Imaging Solution (ThermoFisher Scientific) at 37°C. Before each spectroscopy measurement, the cells were visually inspected by imaging. The axial position for the spectroscopy measurements was placed in the middle of the chosen cell. Multiple planar positions in cells were selected to probe different points (in the cytoplasm or inside SGs) at different temporal points of the process. The fluorescence intensity was acquired for about 120 seconds and analyzed offline. All measurements were performed at 37°C inside a temperature-controlled chamber (Bold Line Temperature Controller, Okolab, PA, USA).

### Image reconstruction and analysis

We reconstructed the ISM images with the adaptive pixel-reassignment method (21). In short, we integrated the 4D data set (*ch, x, y*, ∆*t*, where *ch* is the detector element, *x* and *y* the pixel position, and ∆*t* the tag of the photon arrival time) created in time-tagging modality along the ∆*t* dimension. We applied a phase-correlation registration algorithm to align all the images (*x*, | *y ch*) with respect to the central image. The registration generated the so-called shift-vector fingerprint (*s*_*x*_(*ch*), *s*_*y*_(*ch*)). To obtain the ISM intensity-based images, we integrated the shifted data set along the *ch* dimension. To obtain the lifetime-based ISM image, we started from the 4D data set (*ch, x, y*, ∆*t*); for each ∆*t* value, we used the same shift-vector fingerprint to shift the relative 2D image; we integrated the result along the *ch* dimension; we used the resulting 3D data-set (*x, y*, ∆*t*) and the plugin FLIMJ (based on ImageJ) (48) to obtain the *τ*_*fl*_ maps (fitted with a single-exponential decay model).

Alternatively, we applied the phasor analysis on the same 3D data set. Phasor analysis allows us to interpret the fluorescence lifetime by projecting the photon arrival-time histograms in a 2D coordinate system without the need for fitting. We calculated the phasor coordinates (*g, s*) using cosine and sine summations (49, 50). To avoid artifacts, we performed the MOD mathematical operation of the time-correlated single photon counting histograms with the laser repetition period value (50). Importantly, because each SPAD element can introduce a specific, but fixed, delay, we calibrated the system by measuring the instrument response function of the complete setup (microscope, detector, and BrightEyes-TTM) with a solution of Atto-495 (ATTO-TEC GmbH Germany, *τ*_fl_ = 1 ns), or of Atto-594 (ATTO-TEC GmbH, *τ*_fl_ = 3.9 ns).

### Fluorescence correlation calculation and analysis

We calculated the correlations directly on the lists of absolute photon times (51). For the sum 3 ×3 and sum 5× 5 analysis, the lists of all relevant SPAD channels were merged and the correlations were calculated. The data was then split into chunks of 10 or 5 s, depending on the probe observed, and for each chunk, the correlations were calculated. The individual correlation curves were visually inspected, and all curves without artifacts were averaged. For both single-point and circular scanning (52) FCS, the correlation curves were fitted with a 1-component model assuming a Gaussian detection volume, as described in (6). The theoretical fitting functions for FCS are briefly summarized in the Supplementary Information 1. For the circular FCS measurements (52), the periodicity and radius of the scan movement were kept fixed while the amplitude, diffusion time, and focal spot size were fitted. This procedure was used for the fluorescent beads and allowed calibrating the different focal spot sizes (i.e., central, sum 3 ×3 and sum 5× 5). For the conventional FCS measurements, the focal spot size was kept fixed at the values found with circular scanning FCS, and the amplitude and diffusion times were fitted. Since we approximated the PSF as a 3D Gaussian function with a 1/*e*^2^ lateral radius of *ω*_0_ and a 1/*e*^2^ height of *k × ω*_0_ (with *k* the eccentricity of the detection volume, *k* = 4.5 for the central element probing volume, *k* = 4.1 for the sum 3 ×3 and sum 5 ×5 probing volume), the diffusion coefficient *D* can be calculated from the diffusion time *τ*_*D*_ and the focal spot size *ω*_0_ via 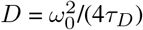

## Data availability

A comprehensive demo dataset of an FFS experiment and the relative Jupyter notebooks for analysis will be available for download. Due to the very large amount of data and diversity, not all data described in the manuscript are currently available to download but can be obtained from the corresponding author upon reasonable request.

## Code availability

The source code for analyzing the photon time-tagging measurements (fluorescence lifetime fluctuation spectroscopy and fluorescence lifetime image scanning microscopy) has been deposited in the GitHub BrightEyes-TTM repository as a part of a larger open-hardware/software project aims at democratising single-photon laser-scanning microscopy based on SPAD array detector. The source code for analyzing the intensity-based measurements (fluorescence fluctuation spectroscopy) will be deposited on the GitHub BrightEyes-FFS repository.

## AUTHOR CONTRIBUTIONS

G.V. conceived the idea. E.P., E.S. and G.V. developed the methodologies for the study. E.Z., G.G.T., I.B., and G.V. supervised and coordinated the project. F.C., D.M., and E.V. prepared the living cell sample. S.Z., J.R., and E.Z. prepared the *invitro* sample. E.P., S.Z., and F.C. performed the living cell experiments. E.P., S.Z., and J-R. performed the *in-vitro* experiments. E.P., S.Z., and G.V. analyzed the living cell and *in-vitro* experiments. E.S. and G.V. designed and built, with the help of E.P., the custom laser-scanning microscope. E.S. designed and implemented the microscope control software. E.P. and E.S. designed and implemented the data analysis software. All authors discussed the results. E.P. and G.V. wrote the manuscript with input from all authors.

## ACKNOWLEDGEMENTS

This project has received funding from: the Fondazione San Paolo, “Observation of bio-molecular processes in live-cell with nano-camera”, No. EPFD0098 (E.S., S,Z., and G.V.); the European Research Council, “BrightEyes”, ERC-CoG No. 818699 (G.V.), “ASTRA”, ERC-SyG No. 855923 (G.G.T., and I.B.); the European Innovation Council, “ivBM-4PAP”, Pathfinder No. 101098989 (G.G.T.); the European Union’s Horizon 2020 research and innovation program under the Marie Sklodowska-Curie grant agreements, “SMSPAD”, No. 890923 (E.S.) and “MINDED” No. 754490 (E.Z.); the NextGeneration EU PNRR MUR - M4C2 – Action 1.4 - Call “Potenziamento strutture di ricerca e creazione di “campioni nazionali di R&S” (CUP J33C22001130001), “National Center for Gene Therapy and Drugs based on RNA Technology”, No. CN00000041 (G.G.T., I.B, and G.V.); by Associazione Italiana per la Ricerca Sul Cancro, “Circular RNAs: novel players and biomarkers in tumorigenesis”, IG 2019 No. 23053 (I.B.); by Ministero dell’Istruzione, dell’Università e Della ricerca (MIUR), “Non-coding RNAs, new players in gene expression regulation: studying their role in neuronal differentiation and in neurodegeneration”, PRIN2017 No. 2017P352Z4 (I.B.). We thank: all members of the Molecular Microscopy and Spectroscopy group and the RNA Systems Biology group at the Istituto Italiano di Tecnologia (IIT) for the daily discussion and suggestions about the project; Prof. Alberto Diaspro and Dr Paolo Bianchini (Nanoscopy & NIC@IIT, IIT) for valuable discussions; Dr Michele Oneto (Nikon Imaging Center, IIT) for support on the experiments; Prof. Alberto Tosi, Prof. Federica Villa, Dr. Mauro Buttafava (Politecnico di Milano), Dr Marco Castello, and Dr Simonluca Piazza (Genoa Instruments) for the realization of the single-photon-avalanche-diode detector array; all members of the RNA Initiative at the Istituto Italiano di Tecnologia, Prof. Stefano Gustincich (Non-coding RNAs and RNA-based therapeutics, IIT) and Dr Francesco Nicassio (Genomics Science, IIT) for their contribution to the long-term vision of this project.

## Bibliography

1. Haustein, E. & Schwille, P. Fluorescence correlation spectroscopy: Novel variations of an established technique. Annu. Rev. Biophys. Biomol. Struct. 36, 151–169 (2007).

2. Digman, M. A. & Gratton, E. Lessons in fluctuation correlation spectroscopy. Annu. Rev. Phys. Chem. 62, 645–668 (2011).

3. Yu, L. et al. A comprehensive review of fluorescence correlation spectroscopy. Front. Phys. 9 (2021).

4. Scipioni, L., Lanzanó, L., Diaspro, A. & Gratton, E. Comprehensive correlation analysis for super-resolution dynamic fingerprinting of cellular compartments using the zeiss airyscan detector. Nat. Commun. 9 (2018).

5. Slenders, E. et al. Confocal-based fluorescence fluctuation spectroscopy with a SPAD array detector. Light Sci. Appl. 10 (2021).

6. Slenders, E. et al. Cooled SPAD array detector for low light-dose fluorescence laser scanning microscopy. Biophys. Rep. 1, 100025 (2021).

7. Wawrezinieck, L., Rigneault, H., Marguet, D. & Lenne, P.-F. Fluorescence correlation spectroscopy diffusion laws to probe the submicron cell membrane organization. Biophys. J. 89, 4029 – 4042 (2005).

8. Ruprecht, V., Wieser, S., Marguet, D. &Schütz, G. J. Spot variation fluorescence correlation spectroscopy allows for superresolution chronoscopy of confinement times in membranes. Biophys. J. 100, 2839–45 (2011).

9. Eggeling, C. et al. Direct observation of the nanoscale dynamics of membrane lipids in a living cell. Nature 457, 1159–1162 (2009).

10. Vicidomini, G. et al. Spatio-temporal heterogeneity of lipid membrane dynamics revealed by STED-FLCS. Nano Lett. 15, 5916–5918 (2015).

11. Lanzanò, L. et al. Measurement of nanoscale three-dimensional diffusion in the interior of living cells by sted-fcs. Nat. Commun. 8, 65 (2017).

12. Bag, N., Ng, X. W., Sankaran, J. & Wohland, T. Spatiotemporal mapping of diffusion dynamics and organization in plasma membranes. Methods Appl. Fluoresc. 4, 034003 (2016).

13. Sankaran, J. & Wohland, T. Fluorescence strategies for mapping cell membrane dynamics and structures. APL Bioeng. 4, 020901 (2020).

14. Huff, J. The airyscan detector from ZEISS: confocal imaging with improved signal-to-noise ratio and super-resolution. Nat. Methods 12, i–ii (2015).

15. Antolovic, I. M., Bruschini, C. & Charbon, E. Dynamic range extension for photon counting arrays. Opt. Express 26, 22234 (2018).

16. Buttafava, M. et al. SPAD-based asynchronous-readout array detectors for image-scanning microscopy. Optica 7, 755 (2020).

17. Rossetta, A. et al. The BrightEyes-TTM as an open-source time-tagging module for democratising single-photon microscopy. Nat. Commun. 13 (2022).

18. Becker, W. Fluorescence lifetime imaging - techniques and applications. J. Microsc. 247, 119–136 (2012).

19. Wallrabe, H. & Periasamy, A. Imaging protein molecules using FRET and FLIM microscopy. Curr. Opin. Biotechnol. 16, 19–27 (2005).

20. Schwille, P., Meyer-Almes, F. & Rigler, R. Dual-color fluorescence cross-correlation spectroscopy for multicomponent diffusional analysis in solution. Biophys. J. 72, 1878–1886 (1997).

21. Castello, M. et al. A robust and versatile platform for image scanning microscopy enabling super-resolution FLIM. Nat. Methods 16, 175–178 (2019).

22. Koho, S. V. et al. Two-photon image-scanning microscopy with SPAD array and blind image reconstruction. Biomed. Opt. Express 11, 2905 (2020).

23. Glauninger, H., Hickernell, C. J. W., Bard, J. A. & Drummond, D. A. Stressful steps: Progress and challenges in understanding stress-induced mRNA condensation and accumulation in stress granules. Mol. Cell 82, 2544–2556 (2022).

24. Guo, Q., Shi, X. & Wang, X. RNA and liquid-liquid phase separation. Non-coding RNA Res. 6, 92–99 (2021).

25. André, A. A. M. & Spruijt, E. Liquid–liquid phase separation in crowded environments. Int. J. Mol. Sci. 21, 5908 (2020).

26. Protter, D. S. & Parker, R. Principles and properties of stress granules. Trends Cell Biol. 26, 668–679 (2016).

27. Patel, A. et al. A liquid-to-solid phase transition of the ALS protein FUS accelerated by disease mutation. Cell 162, 1066–1077 (2015).

28. Bosco, D. A. et al. Mutant FUS proteins that cause amyotrophic lateral sclerosis incorporate into stress granules. Hum. Mol. Genet. 19, 4160–4175 (2010).

29. Mouttou, A. et al. Quantifying membrane binding and diffusion with fluorescence correlation spectroscopy diffusion laws. Biophys. J. (2023).

30. Goedert, M. Alpha-synuclein and neurodegenerative diseases. Nat. Rev. Neurosci. 2, 492– 501 (2001).

31. Ray, S. et al. Alpha-synuclein aggregation nucleates through liquid–liquid phase separation. Nat. Chem. 12, 705–716 (2020).

32. Nath, S., Meuvis, J., Hendrix, J., Carl, S. A. & Engelborghs, Y. Early aggregation steps in alpha-synuclein as measured by FCS and FRET: Evidence for a contagious conformational change. Biophys. J. 98, 1302–1311 (2010).

33. Kroschwald, S., Maharana, S. & Simon, A. Hexanediol: a chemical probe to investigate the material properties of membrane-less compartments. Matters (2017).

34. Dada, S. T. et al. Spontaneous nucleation and fast aggregate-dependent proliferation of alpha-synuclein aggregates within liquid condensates at neutral pH. Proc. Natl. Acad. Sci. U. S. A. 120 (2023).

35. Vance, C. et al. Mutations in FUS, an RNA processing protein, cause familial amyotrophic lateral sclerosis type 6. Science 323, 1208–1211 (2009).

36. Kwiatkowski, T. J. et al. Mutations in the fus/tls gene on chromosome 16 cause familial amyotrophic lateral sclerosis. Science 323, 1205–1208 (2009).

37. Morlando, M. et al. FUS stimulates microRNA biogenesis by facilitating co-transcriptional drosha recruitment. EMBO J. 31, 4502–4510 (2012).

38. Wieser, S., Moertelmaier, M., Fuertbauer, E., Stockinger, H. &Schütz, G. J. (un)confined diffusion of CD59 in the plasma membrane determined by high-resolution single molecule microscopy. Biophys. J. 92, 3719–3728 (2007).

39. Tittelmeier, J., Druffel-Augustin, S., Alik, A., Melki, R. & Nussbaum-Krammer, C. Dissecting aggregation and seeding dynamics of α-syn polymorphs using the phasor approach to FLIM. Commun. Biol. 5 (2022).

40. Sarkisyan, K. et al. Green fluorescent protein with anionic tryptophan-based chromophore and long fluorescence lifetime. Biophys. J. 109, 380–389 (2015).

41. Kim, C., Kang, D., Lee, E. K. & Lee, J.-S. Long noncoding RNAs and RNA-binding proteins in oxidative stress, cellular senescence, and age-related diseases. Oxid. Med. Cell. Longevity 2017, 1–21 (2017).

42. Campos-Melo, D., Hawley, Z. C. E., Droppelmann, C. A. & Strong, M. J. The integral role of RNA in stress granule formation and function. Front. Cell Dev. Biol. 9 (2021).

43. Luo, J. et al. LncRNAs: Architectural scaffolds or more potential roles in phase separation. Front. Genet. 12 (2021).

44. Mitrea, D. M., Mittasch, M., Gomes, B. F., Klein, I. A. & Murcko, M. A. Modulating biomolecular condensates: a novel approach to drug discovery. Nat. Rev. Drug Discovery 21, 841–862 (2022).

45. Castello, M. et al. Universal removal of anti-stokes emission background in STED microscopy via FPGA-based synchronous detection. Rev. Sci. Instrum. 88, 053701 (2017).

46. Rupert, J., Monti, M., Zacco, E. & Tartaglia, G. G. A computational approach reveals the ability of amyloids to sequester RNA: the alpha synuclein case (2022).

47. Martone, J. et al. SMaRT lncRNA controls translation of a g-quadruplex-containing mRNA antagonizing the DHX36 helicase. EMBO reports 21 (2020).

48. Gao, D. et al. FLIMJ: An open-source ImageJ toolkit for fluorescence lifetime image data analysis. PLOS One 15, e0238327 (2020).

49. Digman, M. A., Caiolfa, V. R., Zamai, M. & Gratton, E. The phasor approach to fluorescence lifetime imaging analysis. Biophys. J. 94, L14–L16 (2008).

50. Ranjit, S., Malacrida, L., Jameson, D. M. & Gratton, E. Fit-free analysis of fluorescence lifetime imaging data using the phasor approach. Nat. Protoc. 13, 1979–2004 (2018).

51. Wahl, M., Gregor, I., Patting, M. & Enderlein, J. Fast calculation of fluorescence correlation data with asynchronous time-correlated single-photon counting. Opt. Express 11, 3583 (2003).

52. Petrášek, Z., Derenko, S. & Schwille, P. Circular scanning fluorescence correlation spectroscopy on membranes. Opt. Express 19, 25006 (2011).

