## Supplementary File for "Content-enriched fluorescence lifetime fluctuation spectroscopy to study bio-molecular condensate formation"

### Supplementary Note 1: FCS theory

Fluorescence Correlation Spectroscopy (FCS) is a statistical correlation method based on the detected single-molecule fluctuation of fluorescence intensity which can be used to describe the movement of fluorescently tagged molecules. It was the first method, within the bigger family of Fluorescence Fluctuation Spectroscopy (FFS) to be experimental and theoretical developed, and it represents the baseline method for all the subsequent implementations. Here, there will be a brief step-wise mathematical derivation of the autocorrelation function (ACF). The normalized ACF is defined as follows:

$$G(\tau) = \frac{\langle \delta F(t) \cdot \delta F(t+\tau) \rangle}{\langle F(t) \rangle^2}, \tag{S1}$$

where $\delta F(t)$ represents the fluctuation of fluorescence intensity in a certain time $t$ measured on the detector, and $\tau$, called lag-time, is the time delay relative to an earlier time point in the measurement. The fluctuations of the fluorescence intensity recorded from the detector can be written as:

$$\delta F(t) = k \int_V I_{exc}(\vec{r}) S(\vec{r}) \delta(\sigma_{abs} \Phi C((\vec{r}), t)) dV, \tag{S2}$$

with $S(\vec{r})$ as the optical transfer function of the objective-pinhole, $V$ the volume, $C$ the concentration of the diffusing fluorophores, $\sigma_{abs}$ the cross-section for the absorption process, $k$ the quantum efficiency for the photon detection and $\Phi$ the quantum yield for the emission. If we consider the spatial distribution of the emitted light on the detector plane:

$$W(\vec{r}) = I_{exc}(\vec{r}) S(\vec{r}) \tag{S3}$$

and the molecular brightness, defined as:

$$b = k \sigma_{abs} \Phi \tag{S4}$$

we can simplify equation S2 to:

$$\delta F(t) = \int_V W(\vec{r}) \delta(bC((\vec{r}), t)) dV. \tag{S5}$$

Considering a fluorophore with a constant brightness over time, thus, no photo-physical effects, only the concentration changes are contributing to the detected fluctuation of fluorescence intensity ($\delta(bC(t)) = b\delta C(t)$). In this case, we can re-write equation S1 as:

$$G(\tau) = \frac{\iint_{VV'} W(\vec{r}) W(\vec{r'}) \left\langle \delta C(\vec{r}, 0) \delta C(\vec{r'}, \tau)) \right\rangle dV dV'}{(\int_V \delta C(\vec{r}, 0) W(\vec{r}) dV)^2} \tag{S6}$$

with $t = 0$ for simplicity. If the fluorescently tagged molecules are freely diffusing in a three-dimensional space, the fluctuation of fluorescence given by the changes in concentrations can be described by taking advantage of the solution of the Brownian motion equation:

$$\left\langle \delta C(\vec{r}, 0) \delta C(\vec{r'}, \tau)) \right\rangle = \langle C \rangle \frac{1}{(4\pi D \tau)^{3/2}} e^{(\vec{r} - \vec{r'})^2 / 4D\tau}, \tag{S7}$$

where $D$ is the diffusion coefficient. In the confocal microscopy case, we can describe the detection plane explicitly:

$$W(\vec{r}) = I_0 e^{-2(x^2+y^2)/\omega_0^2} e^{-2z^2/z_0^2}, \tag{S8}$$

with $\vec{r} = (x, y, z)$, and $\omega_0$ and $z_0$ the beam profile parameters. We can introduce the effective volume $V_{eff}$ as:

$$V_{eff} = \frac{(\int_V W(\vec{r}) dV)^2}{\int_V W(\vec{r})^2 dV}, \tag{S9}$$

which is equal to $V_{eff} = (\pi/2)^{3/2} \omega_0^2 z_0$ in the confocal case. Evaluating equation S6 with equations S7 and S9, we can explicitly write the ACF for fluorescently tagged molecules diffusing with Brownian motion in the effective volume defined by the Point Spread Function (PSF):

$$G_D(\tau) = \frac{0.35}{\langle C \rangle V_{eff}} \frac{1}{1 + \frac{\tau}{\tau_D}} \frac{1}{\sqrt{1 + \frac{\tau}{\tau_D} (\omega_0 / z_0)^2}}, \tag{S10}$$

with $\tau_D$ being the diffusion time, defined as $\tau_D = \omega_0^2 / 4D$. The average number of molecules in the effective volume can be calculated from the average concentration as:

$$N = \langle C \rangle V_{eff}, \tag{S11}$$

while the radius of the molecule is related to its diffusion coefficient in the spherical-geometry case.

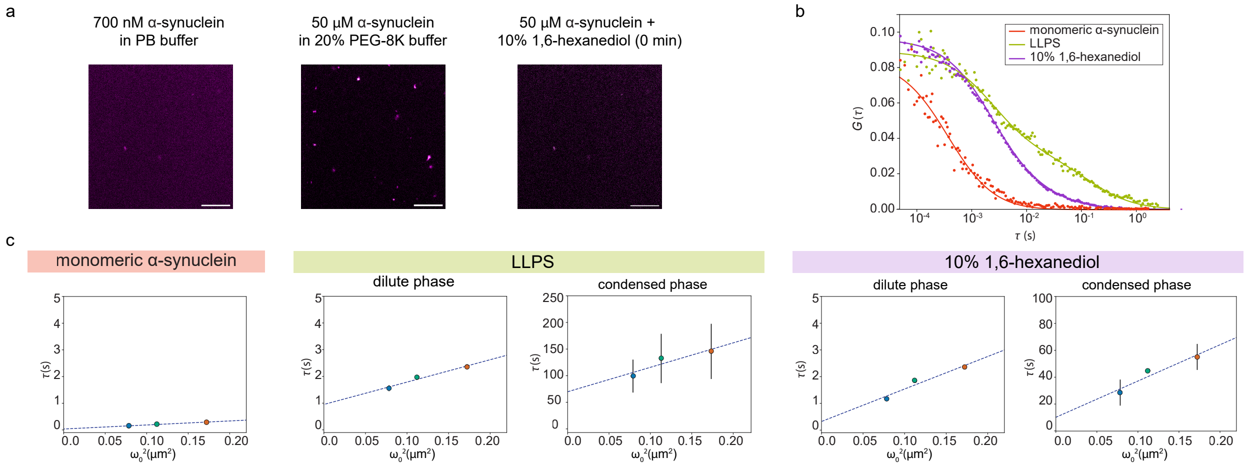


**Fig. S1. Alpha-synuclein LLPS *in vitro*.** **a** ISM intensity-based images of 700 nM alpha-synuclein ($\alpha$-syn) before LLPS (20 mM phosphate buffer (PB), *left*) and 50 µM $\alpha$-syn (5% $\alpha$-syn$^{C141}$-atto488) in LLPS buffer (20% PEG-8K, 20 mM phosphate buffer, *middle*) and immediately after the addition of 10% 1,6-hexanediol (20% PEG-8K, 20 mM PB, *right*). Scale bars 10 µm. **b** Autocorrelation curves of $\alpha$-syn in the three conditions depicted in (a). Red for $\alpha$-syn in 20 mM PB, green for $\alpha$-syn in LLPS buffer, and purple for dissolved $\alpha$-syn condensates. **c** Diffusion law plots for the three conditions of $\alpha$-syn condensation. While $\alpha$-syn in 20 mM PB can be analyzed with one diffusing component only, $\alpha$-syn after the LLPS and after the addition of 10% 1,6-hexanediol are analyzed with a two components FCS model. In both cases, the first component represents the diluted phase, having an intercept close to zero ($q$ = (0.9$\pm$0.1) for LLPS $\alpha$-syn and $q$ = (0.3$\pm$0.3) for 10% 1,6-hexanediol $\alpha$-syn). The second component represents the condensed phase (intercept bigger than zero, $q$ = (70$\pm$20) and $D$ = (0.5$\pm$0.1) µm$^2$/s) for $\alpha$-syn in LLPS, and some partial not-dissolved smaller condensates ($q$ = (13$\pm$2) and $D$ = (0.8$\pm$0.2) µm$^2$/s) for $\alpha$-syn with %10 1,6-hexanediol.

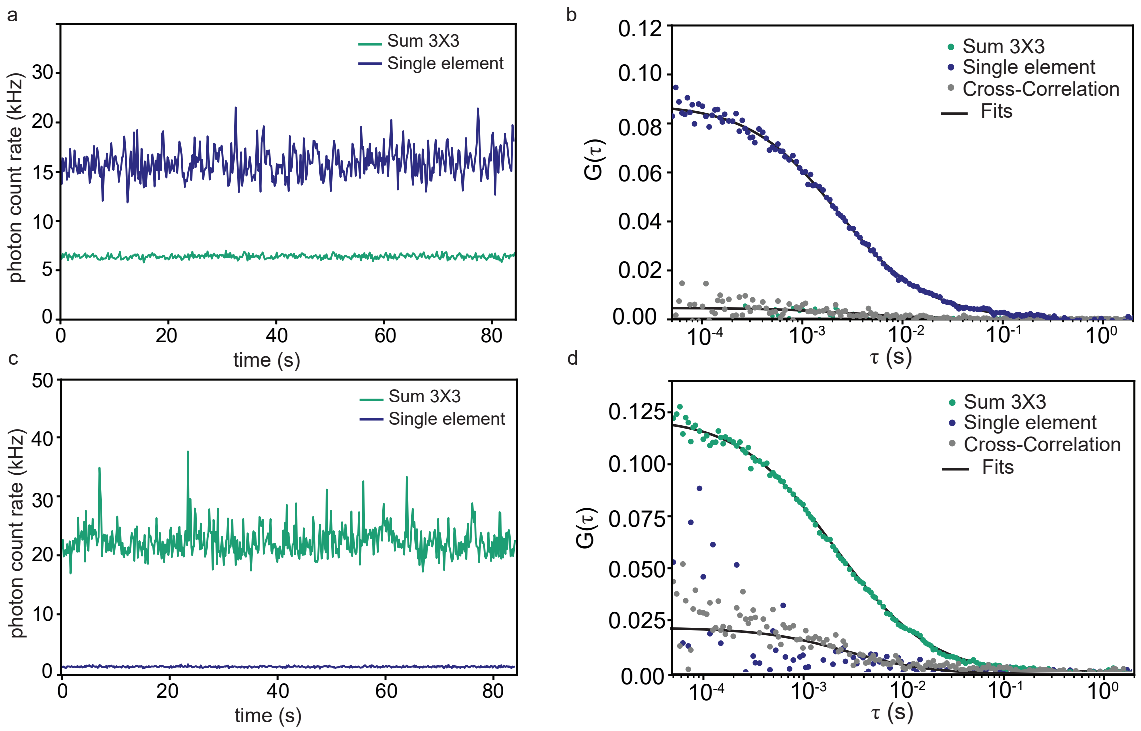


**Fig. S2. Fluorescence Cross-Correlation experiment with orange or yellow-green fluorescent beads only. a** Intensity time trace measured by the two detector arms for a sample of only orange fluorescent beads. **b** Dual-color Fluorescence Cross-Correlation Spectroscopy measurement of red fluorescent nanospheres in water. There is no fluorescence signal from the sample on the SPAD array detector equipped with the green filter; thus, the autocorrelation is null. On the other hand, the single-element detector, equipped with the orange filter, can detect the bead's signal; thus, an autocorrelation curve is calculated. As expected, the cross-correlation between the two is low. **c** Intensity time trace measured by the two detector arms for a sample of only yellow-green fluorescent beads. **d** Dual-color Fluorescence Cross-Correlation Spectroscopy measurement of yellow-green fluorescent nanospheres in water. There is no fluorescence signal from the sample on the single-element detector equipped with the red filter; thus, the autocorrelation is null. On the other hand, the SPAD array detector, equipped with the green filter, can detect the bead's signal; thus, an autocorrelation curve is calculated. As expected, the cross-correlation between the two is low.

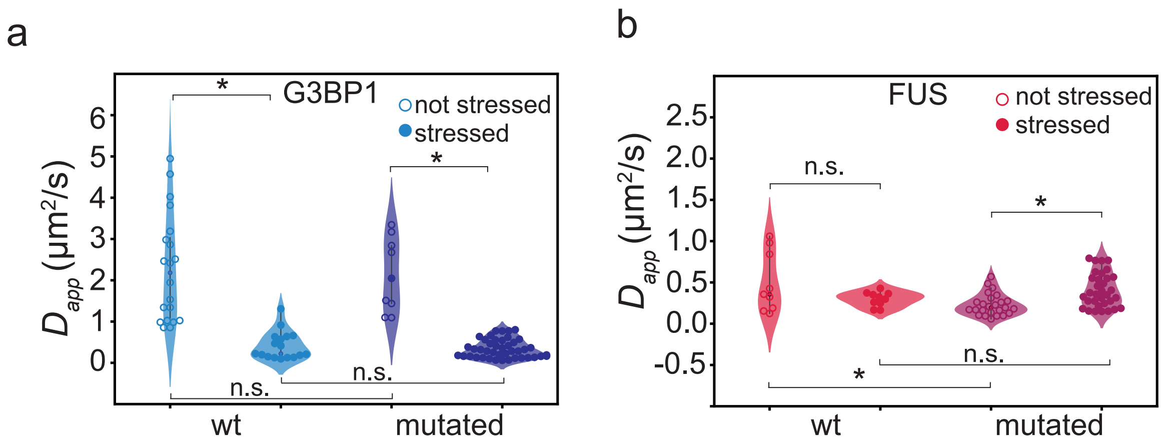


**Fig. S3. G3BP1 and FUS diffusion coefficients. a** Apparent diffusion coefficients of G3BP1 protein before the stress, in the cytosolic phase (empty symbols), and after the stress, in the condensate phase (filled symbols), in the wt (left) and in the mutated condition (right). **b** Apparent diffusion coefficients of FUS protein before the stress, in the cytosolic phase (empty symbols), and after the stress, in the condensate phase (filled symbols), in the wt (left), and in the mutated condition (right). The bars represent the variance of the dataset

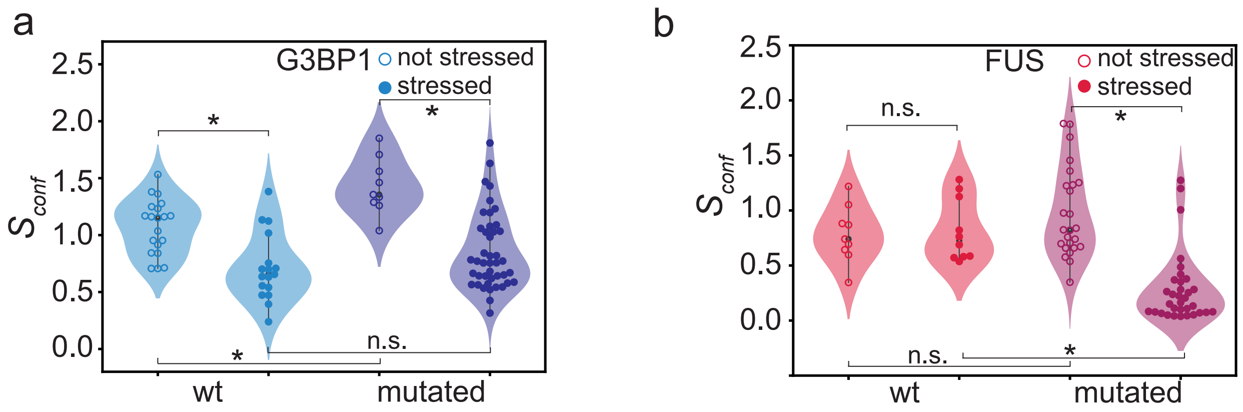


**Fig. S4. G3BP1 and FUS confinement strength a** Confinement strength of G3BP1 protein before the stress, in the cytosolic phase (empty symbols), and after the stress, in the condensate phase (filled symbols), in the wt (left) and in the mutated condition (right). **b** Confinement strength of FUS protein before the stress, in the cytosolic phase (empty symbols), and after the stress, in the condensate phase (filled symbols), in the wt (left), and in the mutated condition (right). The bars represent the variance of the dataset

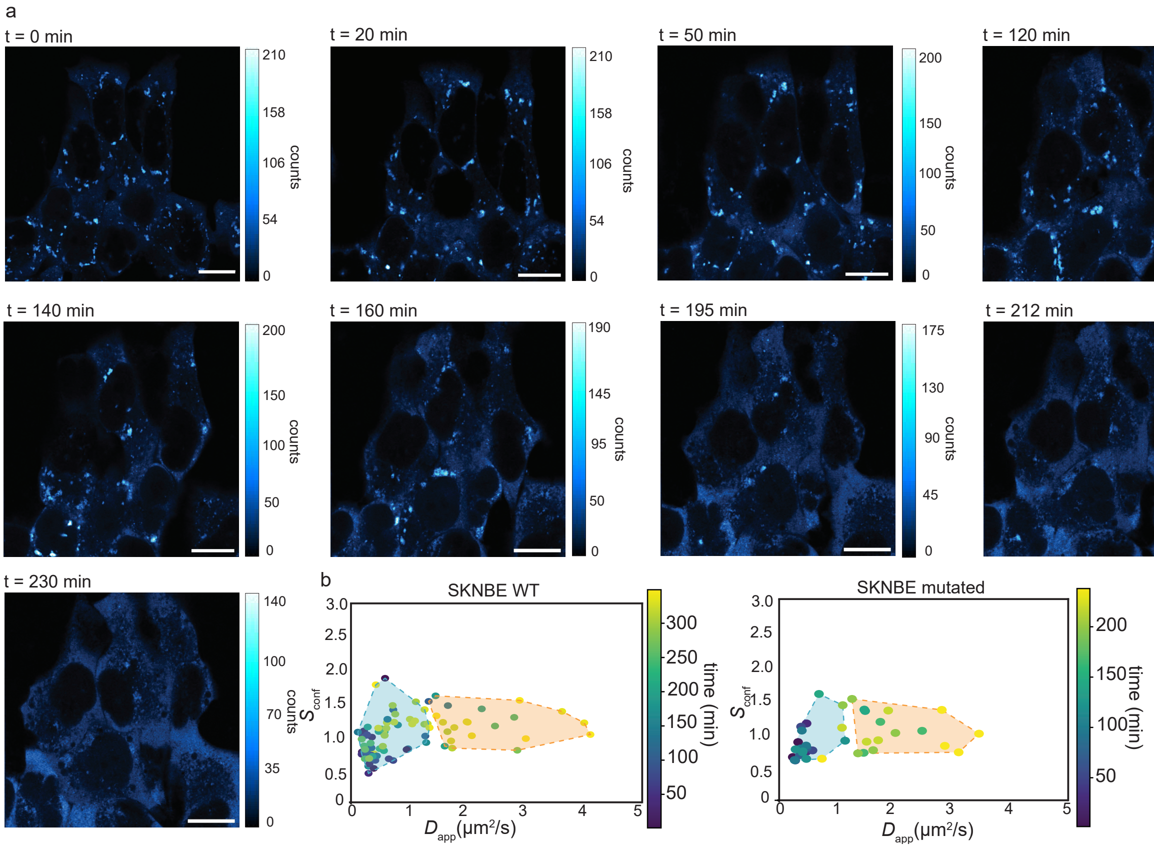


**Fig. S5. Spot-Variation FCS to study stress granule recovery a** Time-lapse ISM imaging of the stress granules recovery. SK-N-BE cells pathologically mutated, expressing G3BP1-gfp and FUS-RFP (here, only G3BP1 is shown) during stress granule recovery. Once the sodium-arsenite is removed, the stress granules dissolve. Scale bars 10 μm. **b** Confinement strength versus diffusion coefficient for G3BP1-GFP in both wt (left) and mutated (right) cell lines. The color code represents the measurement time for each data point. Over time both the confinement strength and the diffusion coefficient retrieve values comparable to the ones before the stress.

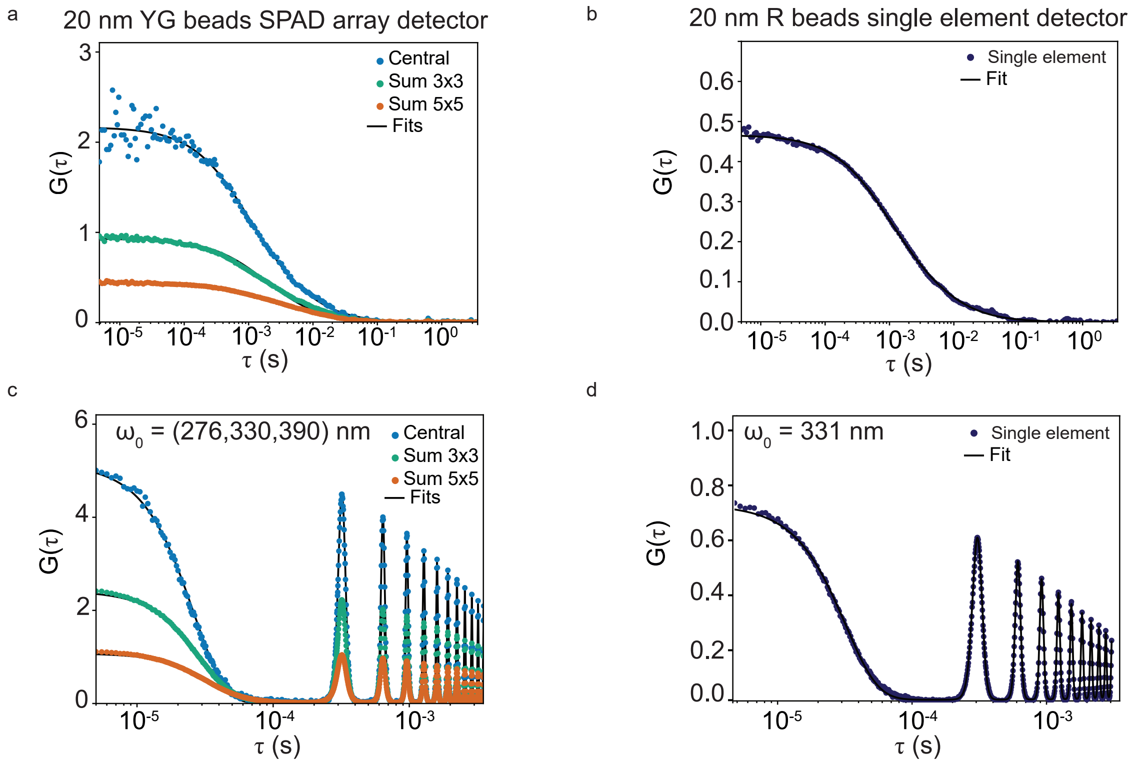


**Fig. S6. Calibration of the detection volumes for FCCS a** Example of a single point FCS measurements of 20 nm YG fluorescent beads with the SPAD array detector. **b** Example of a circular FCS scanning measurements of 20 nm YG fluorescent beads with the SPAD array detector. The detecting volume for this measurement is **c** Example of a single point FCS measurements of 20 nm red fluorescent beads with the SPAD array detector. **d** Example of a circular FCS scanning measurements of 20 nm red fluorescent beads with the SPAD array detector.

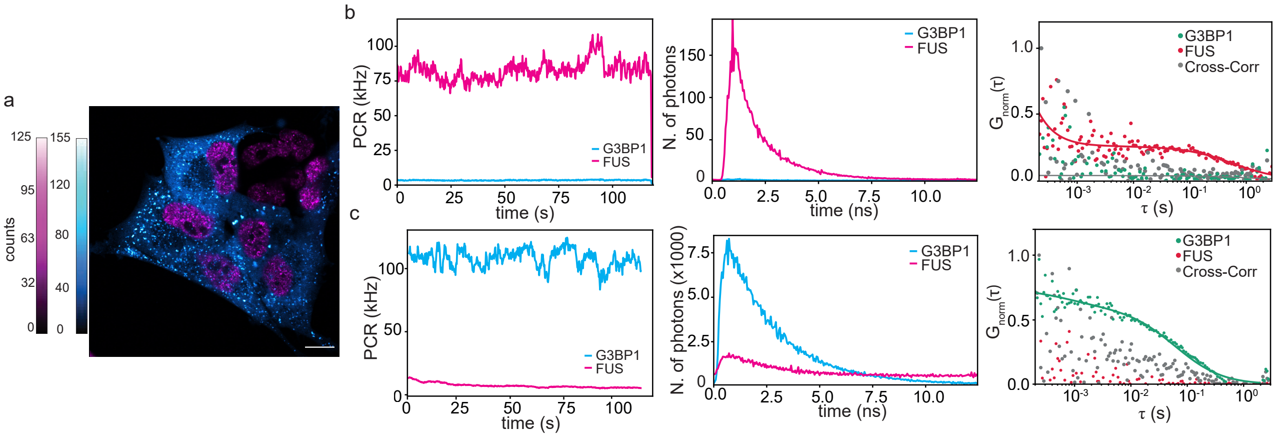


**Fig. S7. FCCS on SK-N-BE wt cells under oxidative stress a** Intensity-based ISM image of a SK-N-BE mutated cell expressing G3BP1-gfp (cytoplasm, in cyan) and FUS (nucleus, magenta) after the oxidative stress. Scale bar 10 µm. **b** FCCS experiment in the nuclei of the cells. Intensity over time (left) of G3BP1-eGFP and FUS-rfp, time decay histograms (center) for G3BP1 (cyan) and FUS (magenta) and autocorrelations and cross-correlation curves (right) (grey) calculated from the same data in the nuclei. **c** FCCS experiment in the cytoplasm of the cells. Intensity over time (left) of G3BP1-eGFP and FUS-rfp, time decay histograms (center) for G3BP1 (cyan) and FUS (magenta) and autocorrelations and cross-correlation curves (right) (grey) calculated from the same data in the cytoplasm.
